# Chromatin accessibility predictions reveal lineage-structured effects of noncoding variants across hematopoiesis

**DOI:** 10.64898/2026.11.12.766027

**Authors:** Mehrdad Kazemi, John A. Morris

## Abstract

**Background:** Noncoding variants account for over 90% of genome-wide association study (GWAS) findings, but their mechanisms are often unclear. Sequence-to-function models provide one approach to variant interpretation by predicting the effects of genetic variation on molecular phenotypes. ChromBPNet predicts variant effects on chromatin accessibility, but how these predictions are organized across differentiation hierarchies and relate to orthogonal predictions remains unclear.

**Results:** We trained ChromBPNet models for 17 hematopoietic cell types and scored 302,830 variants. Of 242,307 variants accessible in at least one cell type, 5.6% were predicted to alter chromatin accessibility. Variants active in more cell types had larger predicted effects. Predicted effects also reflected hematopoietic lineage structure, with sharing declining with lineage distance while effect direction was preserved in 99.6% of parent-child comparisons. ChromBPNet predictions agreed with GWAS fine-mapping but largely identified distinct variants. Fine-mapping-only variants were enriched for coding and transcript-associated variation, whereas ChromBPNet-only variants were enriched for regulatory elements, accessible chromatin, and trait-relevant activity. In K562 CRISPRi datasets, predicted accessibility gain showed limited association with target-gene validation. Regulatory elements containing CTCF motifs were less likely to have validated target genes, consistent with a role in chromatin organization rather than gene activation.

**Conclusions:** ChromBPNet and fine-mapping provide complementary information about variant function. Predicted accessibility effects identify the cell types in which noncoding variants are likely to act and capture relationships across the hematopoietic hierarchy, but provide limited information about the genes they regulate. All predictions are available at: https://bloodnnet.ccbr.utoronto.ca/.

## Background

Most variants associated with blood cell traits lie outside protein-coding sequence, and for most, their molecular consequences, relevant cell types, and target genes remain uncertain [1, 2]. Statistical fine-mapping helps prioritize candidate causal variants but does not establish how those variants function or which genes they affect [3]. Sequence- to-function models provide a complementary approach by predicting the effects of genetic variation on molecular phenotypes in specific cellular contexts [4]. Interpreting these predictions requires distinguishing three related questions: whether a variant alters a molecular phenotype, whether it contributes to a trait association, and which gene mediates its effect.

ChromBPNet is a sequence-based model that predicts chromatin accessibility from DNA sequence while accounting for assay-specific sequence bias [5]. Predicted variant effects are enriched among fine-mapped disease-associated variants and correlate with measured chromatin accessibility QTL effects [5]. ChromBPNet has also been applied across large cell-type atlases to prioritize candidate regulatory variants and interpret genetic associations [6, 7]. However, it remains unclear how predicted accessibility effects are organized across developmental hierarchies, how they complement statistical fine-mapping, and how informative they are for identifying target genes. Addressing these questions is important for understanding both the value and the limitations of sequence-based predictions for noncoding variant interpretation.

Hematopoiesis provides a useful setting in which to investigate these questions. Its branching developmental structure allows predicted accessibility effects to be compared across related cell types [8]. Large-scale fine-mapping results are available for blood cell traits [2], providing an independent source of evidence against which to evaluate model predictions. In addition, CRISPR interference (CRISPRi) screens in K562 cells provide experimental evidence linking candidate cis-regulatory elements (cCREs) to target genes [9–17]. Together, these resources make it possible to evaluate predicted accessibility effects across multiple levels of evidence, from chromatin accessibility to trait association and gene regulation.

Here, we trained ChromBPNet models for 17 hematopoietic cell types (Fig. 1a) and evaluated predicted variant effects using cell-type-specific empirical null distributions. We first characterized how predicted accessibility effects vary across the hematopoietic hierarchy. We then compared model predictions with fine-mapping results for 29 blood cell traits and examined the variants uniquely nominated by each approach. Finally, we evaluated whether predicted accessibility effects distinguish K562 cCREs with experimentally validated target genes.

**Fig. 1.**
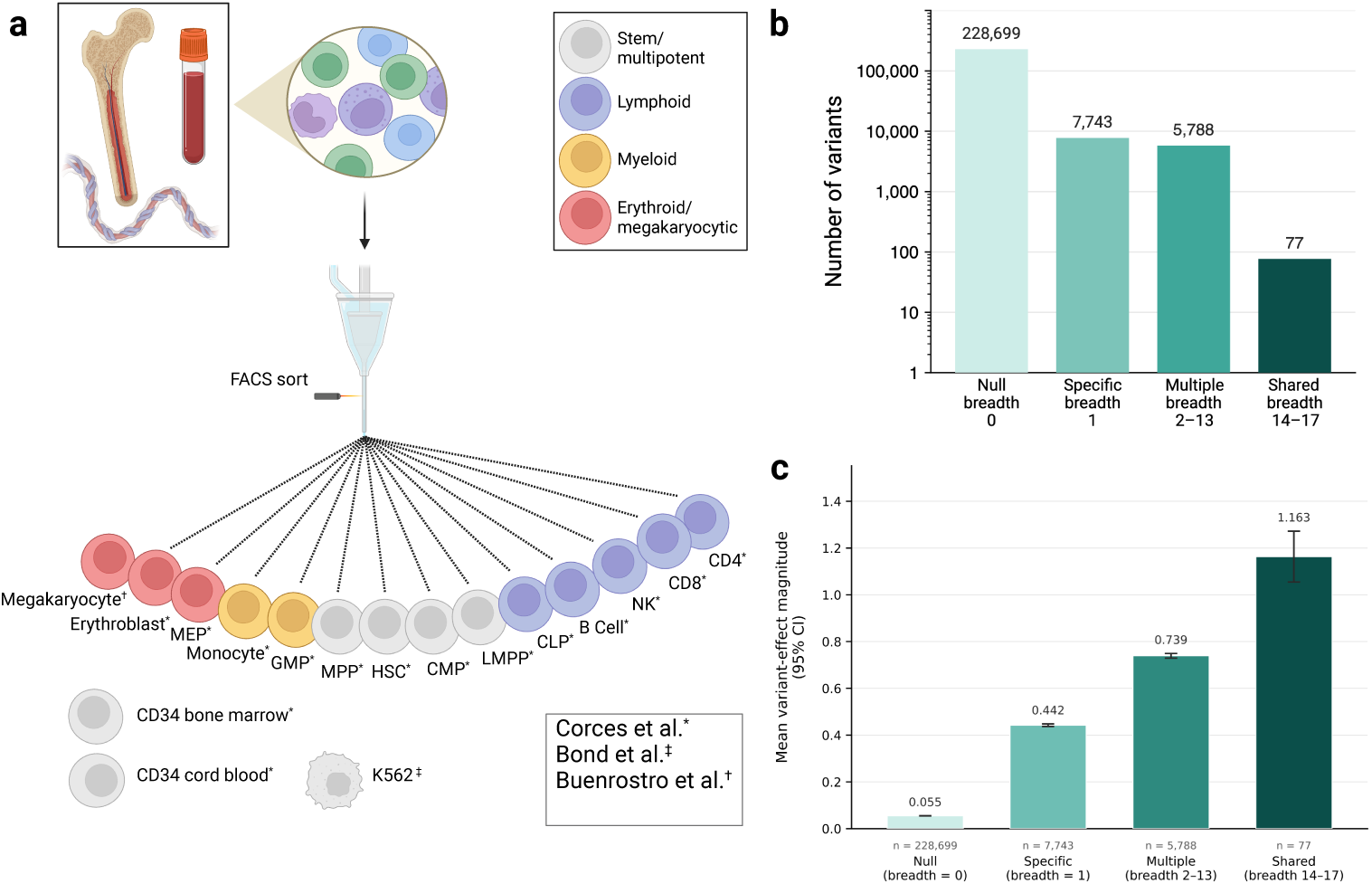
Predicted accessibility effects across the hematopoietic lineage. a, Sources of ATAC- seq data used to train ChromBPNet models for 17 blood cell types. Cells were derived from bone marrow, peripheral blood, and umbilical cord blood. Primary hematopoietic populations from Corces et al. [28] were isolated by FACS, except the CD34^+^ populations. Megakaryocytes and K562 were obtained from separate studies, as indicated by the symbols. Cell types are colored by lineage family (stem/multipotent, lymphoid, myeloid, and erythroid/megakaryocytic). **b,** Number of variants (log scale) in each breadth category, where breadth is the number of the 17 cell types in which a variant is active. A variant was considered active when it was accessible and significant at empirical P *<* 0.01 against that cell type’s shuffled null. Categories are inactive (0 cell types), specific (1), multiple (2- 13), and shared (14-17). **c,** Mean regulatory magnitude by breadth category. Regulatory magnitude was defined as the maximum |logFC| across the cell types in which a variant was accessible. Bars show means and error bars show 95% confidence intervals; category sizes are shown below the x-axis. **a** created in BioRender. Kazemi, M. (2026) https://BioRender.com/u2fjd32.

Predicted accessibility effects followed developmental relationships within the hematopoietic hierarchy and provided information complementary to statistical finemapping. In contrast, they showed limited ability to distinguish cCREs with validated target genes. Together, these results suggest that chromatin accessibility predictions are most informative for identifying where noncoding variants are likely to act, but are less informative for identifying the genes they affect. To support exploration of these predictions, we provide an interactive resource containing model predictions across the hematopoietic lineage.

## Results

### Predicted effects grow with the number of cell types in which a variant is active

We classified 242,307 variants accessible in at least one cell type into four groups based on the number of cell types in which they were predicted to be active (Methods): inactive, active in one cell type (Specific), active in two to thirteen cell types (Multiple), and active in at least fourteen of seventeen cell types (Shared). Most active variants were restricted to a single cell type. Of the 13,608 active variants, 57% were Specific whereas only 77 (0.6%) were Shared (Fig. 1b; Additional file 1: Fig. S1a). Every cell type contributed active variants, with 3.2–4.8% of accessible variants predicted to be active (Additional file 1: Fig. S1b).

Predicted effect size increased with regulatory breadth. We summarized each variant by the largest absolute predicted effect (*|*logFC*|*) observed across accessible cell types and found that mean magnitude increased 2.65-fold from Specific variants (0.44) to Shared variants (1.16) (Fig. 1c). This relationship remained when breadth was defined independently of the effect estimate being tested (Methods; Spearman *ρ* = 0.39, P *<* 10*^−^*^4^) and was absent when breadth was defined using chromatin accessibility rather than predicted activity (*ρ* = -0.02; Additional file 1: Fig. S1c, Note S1). These results are consistent with observations by Marderstein et al. [7], who reported that larger predicted effects tend to occur at variants acting across more cell types. We observed the same relationship within the hematopoietic differentiation hierarchy. Together, these findings suggest that effect size and regulatory breadth capture related but distinct information. Variants should therefore be prioritized using both their predicted effect size and the cell types in which they are active.

We observed the same pattern using AlphaGenome Variant Impact (AVI) [18, 19], a genome-wide prioritization score constructed independently of ChromBPNet. Mean AVI increased from 4.5 in inactive variants to 9.2, 12.7, and 18.3 across the three active breadth classes (Additional file 1: Fig. S1d). Applying a stringent genome-wide threshold (PHRED *≥* 20; top 1% genome-wide) preferentially selected broadly active variants: although single-cell-type variants represented 57% of active variants, only 28% exceeded this threshold. After matching variants for ChromBPNet magnitude and accessibility breadth, broader variants still had higher AVI scores than singlecell-type variants (difference 1.16 units [0.92, 1.40]; Methods; Additional file 1: Note S2). Because AVI is trained to prioritize pathogenicity and evolutionary constraint, this enrichment is expected. However, it also suggests that genome-wide prioritization scores may underprioritize variants acting within a single lineage, even when those variants have strong predicted regulatory effects.

Active variants were also distributed differently across candidate cis-regulatory element (cCRE) classes. Single-cell-type variants were enriched outside annotated cCREs (21% versus 10%; OR 2.4 [2.1, 2.6]), whereas the broadest variants were strongly enriched in promoter-like elements (30% versus 5%; OR 7.7 [4.7, 12.6]) (Methods; Additional file 1: Fig. S2, Note S3). Intermediate-breadth variants were most enriched in distal enhancer-like elements (OR 1.5 [1.4, 1.6]), although distal enhancers were the most common cCRE class across all active categories. Thus, promoter variants tend to act broadly, enhancer variants tend to act across intermediate numbers of cell types, and cCRE annotation alone provides limited information about regulatory breadth.

### Variants active in several cell types are active in lineage-related ones

We next asked whether variants active in multiple cell types tended to act in developmentally related ones. We placed the 14 primary hematopoietic cell types on a differentiation graph and defined lineage distance as the number of edges separating two cell types (Fig. 2a; Methods; Additional file 1: Fig. S3a).

**Fig. 2.**
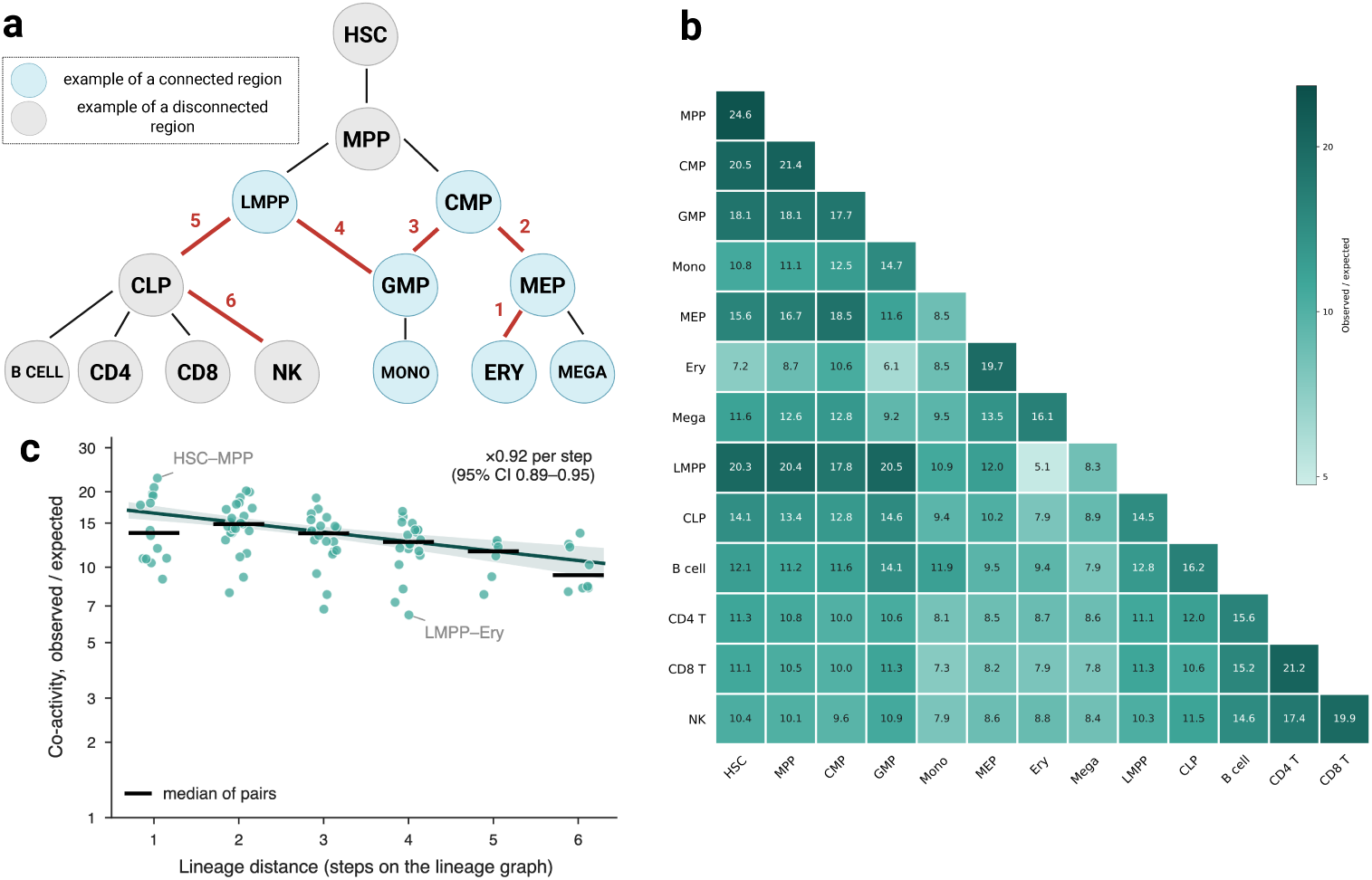
Predicted accessibility effects follow the hematopoietic differentiation hierarchy. a, Lineage graph of the 14 primary hematopoietic cell types used in lineage analyses. Distances between cell types were defined as the number of edges on the shortest path between them. The example shown connects erythroid (ERY) and NK cells by a six-step path (red). **b,** Excess sharing of active variants among all 91 cell-type pairs. Values show the number of variants active in both cell types divided by the number expected under independence. All pairs showed more sharing than expected, with the strongest sharing observed between closely related cell types. **c,** Excess sharing of active variants as a function of lineage distance. Each point represents one of the 91 cell-type pairs. Lineage distance is the shortest-path distance on the lineage graph. The fitted line shows the decline in excess sharing with increasing distance, corresponding to a 0.92-fold decrease per lineage step; the shaded region shows the 95% confidence interval. Black bars indicate the median within each distance class. The y-axis is shown on a logarithmic scale. **a** created in BioRender. Kazemi, M. (2026) https://BioRender.com/u2fjd32.

Variants active in multiple cell types were strongly clustered within the lineage hierarchy. Among the 5,374 variants active in at least two cell types, 47.9% were active in a connected region of the graph compared with 13.5% expected under a permutation null that preserved each variant’s breadth and each cell type’s number of active variants (P *<* 0.001; Methods). The mean distance between active cell types was 2.08 lineage steps compared with 2.89 expected.

Sharing declined with lineage distance. Across all 91 cell-type pairs, variants were more likely to be active in both members when the cell types were developmentally close (Fig. 2b; Additional file 1: Fig. S3b). Each additional lineage step reduced sharing by 8% (*β* = -0.087, 95% CI -0.122 to -0.052; permutation P = 0.001; Fig. 2c; Methods). Nearby cell types also had more similar predicted chromatin effects, with agreement declining from 0.70 between adjacent cell types to 0.46 at six lineage steps (Additional file 1: Fig. S3c). Accounting for this similarity eliminated the association with lineage distance (Additional file 1: Note S4), suggesting that lineage effects are largely captured through similarity in chromatin accessibility.

The direction of effect was almost always preserved across differentiation. Among 11,416 instances where a variant was active in both members of a parent-child celltype pair, 99.6% had the same direction of effect (Methods). This is consistent with a motif-driven mechanism in which sequence changes create or disrupt transcription factor binding sites, while differences in transcription factor activity across cell types modify effect size rather than reverse effect direction.

Together, these results show that regulatory effects are shared preferentially between developmentally related cell types and are generally inherited from progenitor to descendant. Variants predicted to act in one cell type are therefore likely to act in closely related cell types as well.

### ChromBPNet and fine-mapping nominate distinct variant classes

Fine-mapping and ChromBPNet nominate overlapping but largely distinct sets of variants. Among 193,896 GWAS variants with both annotations (Methods), 3,084 were nominated by fine-mapping (PIP *≥* 0.5) and 2,479 by ChromBPNet (active in at least one cell type). Of the 5,324 variants nominated by either method, only 239 were nominated by both (Fig. 3).

**Fig. 3.**
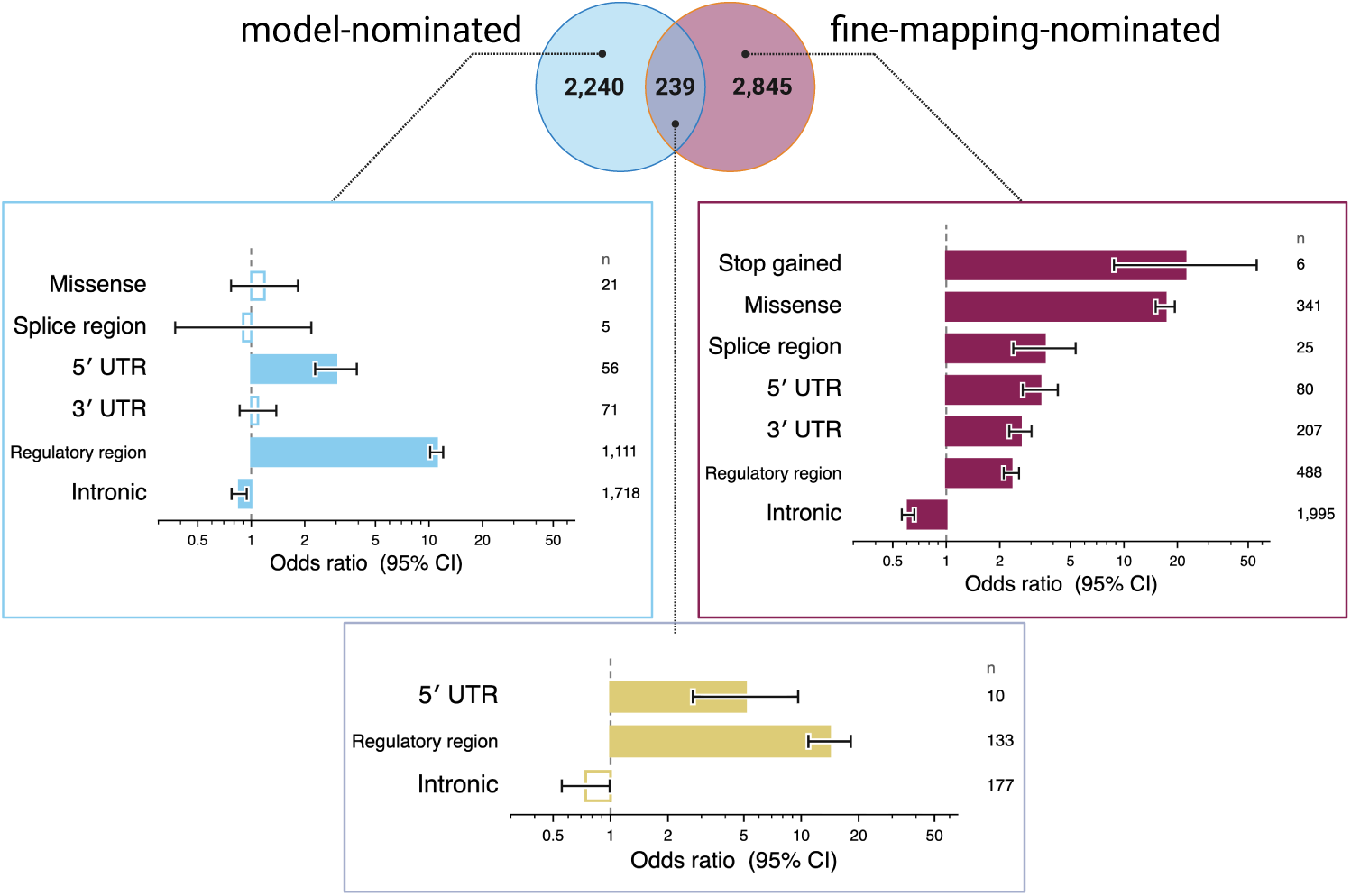
Fine-mapping and ChromBPNet nominate overlapping but distinct classes of variants. The Venn diagram shows overlap between variants nominated by fine-mapping (PIP *≥* 0.5 for at least one of 29 blood cell traits) and ChromBPNet (active in at least one of 17 blood cell types), among the 193,896 variants with both annotations. Variant counts are shown for the fine-mapping-only (n = 2,845), both (n = 239), model-only (n = 2,240), and neither (n = 188,572) classes. For each nominated class, bars show enrichment or depletion of selected Ensembl VEP consequences relative to variants nominated by neither method. Filled bars indicate consequences significant after Benjamini- Hochberg correction (q *<* 0.05); open bars indicate q *≥* 0.05. Error bars show 95% confidence intervals. The seven consequence classes discussed in the text are shown: stop gained, missense, splice region, 5*^′^* UTR, 3*^′^* UTR, regulatory region, and intronic.

The overlap was greater than expected among variants in accessible chromatin. Among the 36,200 accessible variants, fine-mapped variants were four times more likely than other variants to be nominated by ChromBPNet (OR 4.11, 95% CI 3.54-4.77). Agreement increased with fine-mapping confidence. Among variants with PIP *≥* 0.9, 27% were also nominated by ChromBPNet, while 73% were not (Additional file 1: Note S5). The two approaches therefore identify overlapping but distinct groups of variants.

The 88 variants nominated most strongly by both methods represent a highconfidence set for functional follow-up (Methods; Additional file 1: Fig. S4; Additional file 2: Table S1). For example, we identified rs2814778, previously reported as the Duffy-null variant [20], which has a PIP of 1.0 for white cell counts, neutrophil counts and lymphocyte percentage. ChromBPNet predicts reduced accessibility across erythroid and myeloid cell types but not lymphoid cells, consistent with its known erythroid-restricted effect through disruption of a GATA1 motif [20].

We also release annotations for all 13,608 active variants (Additional file 2: Table S2). This set includes rs112233623, a known erythroid regulatory variant in a CCND3 enhancer [21], which has a PIP of 1.0 for ten red cell indices. ChromBPNet predicts reduced accessibility in MEP, erythroid cells, and K562, consistent with its established role in red blood cell biology.

### Fine-mapping-only and ChromBPNet-only variants differ in what they are likely to do

Variants nominated only by fine-mapping and only by ChromBPNet show different functional signatures. Of the 2,845 variants nominated only by fine-mapping, 70% lie outside accessible chromatin and therefore cannot be nominated by a model of chromatin accessibility. Among those in accessible chromatin, fine-mapping-only variants are enriched for missense and 5*^′^* untranslated-region variants and depleted for intronic variants (Fig. 3; Methods; Additional file 1: Fig. S5, Note S6). These are classes of variation expected to act through protein sequence or transcript processing rather than chromatin accessibility.

Variants nominated only by ChromBPNet show the opposite pattern. They are enriched for annotated regulatory elements and show no enrichment for missense variation. Variants nominated by both methods carry the same regulatory signature (Fig. 3; Additional file 1: Fig. S5).

ChromBPNet-only variants also occur in more accessible chromatin than finemapping-only variants. The median active-allele accessibility quantile was 0.692 for ChromBPNet-only variants compared with 0.607 for fine-mapping-only variants (q = 3 *×* 10*^−^*^10^; Methods; Additional file 1: Note S6). ChromBPNet-only variants therefore do not represent large predicted effects in weakly accessible regions.

Together, these results suggest that disagreement between the two methods reflects differences in the variants they prioritize. Fine-mapping-only variants are enriched for coding and transcript-associated variation, whereas ChromBPNet-only variants are enriched for regulatory variation in accessible chromatin.

### Model-only variants act in the cell types of their traits

Cell-type specificity provides an independent test of the model-only variants because neither ChromBPNet nor fine-mapping uses trait-cell-type relationships. We therefore asked whether variants associated with a given blood cell lineage were active in cell types from that lineage.

Model-only variants showed significant lineage concordance. Among 4,667 varianttrait pairs, 39.9% were active in the expected lineage compared with 31.0% under a permutation null (Methods; 1.29-fold; P *<* 10*^−^*^4^). Variants nominated by both methods showed even stronger concordance (73.3% observed versus 45.9% expected; 1.60-fold; P *<* 10*^−^*^4^), consistent with their higher confidence.

Together with their enrichment in regulatory elements and accessible chromatin, these results suggest that many model-only variants are genuine regulatory variants that remain unresolved by fine-mapping.

### Predicted accessibility gain weakly distinguishes CRISPRi-validated cCREs in K562

We next asked whether variants with larger predicted effects on chromatin accessibility are more likely to occur in regulatory elements with validated target genes. We compiled 9,205 cCREs from published K562 CRISPRi screens [9–17], including 1,259 with at least one validated target gene, and scored blood trait GWAS [2] and blood eQTL variants [22] that fell within them (Fig. 4a; Methods). Of these, 2,801 cCREs contained at least one scored variant.

**Fig. 4.**
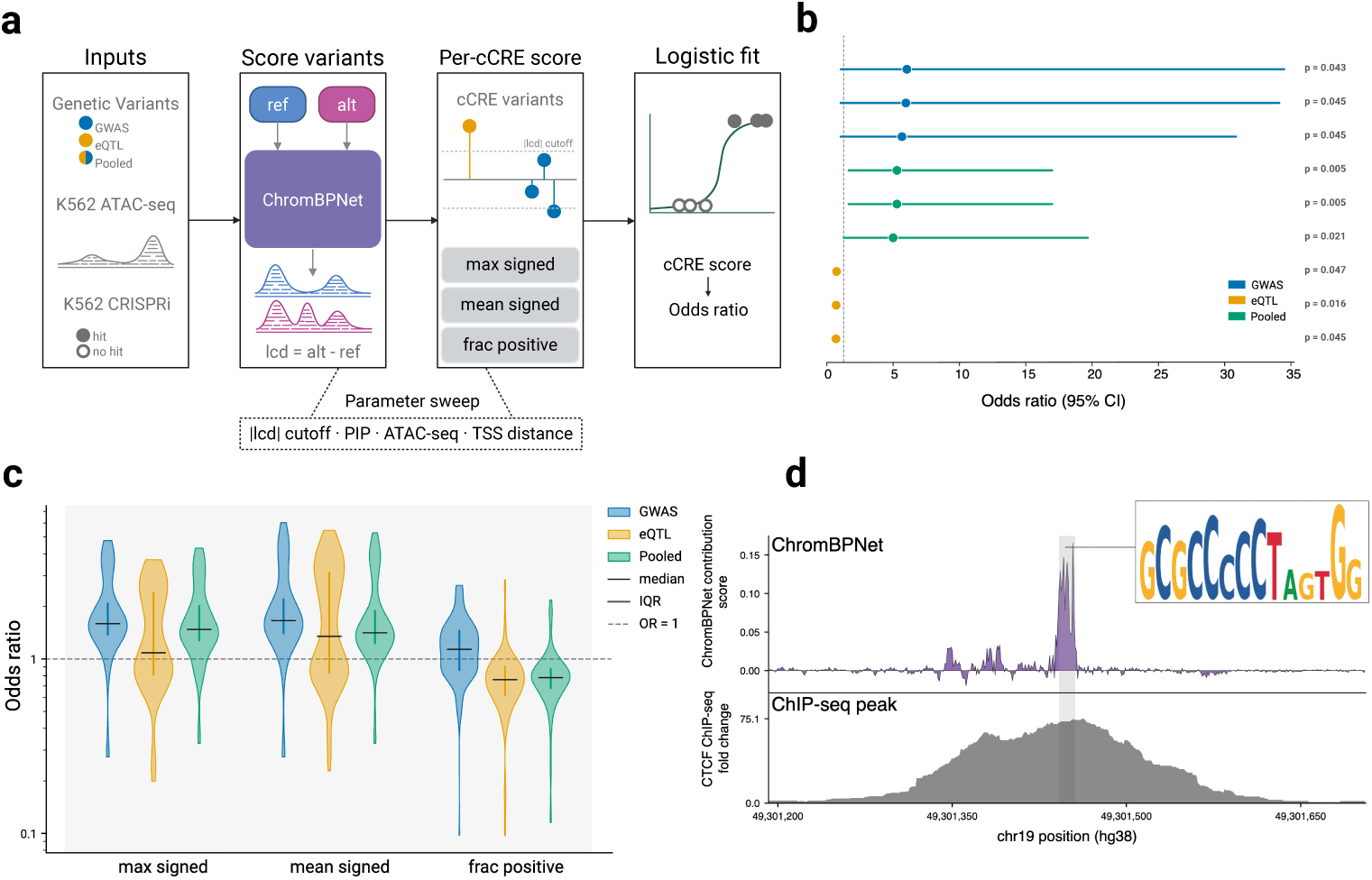
ChromBPNet predictions show limited ability to identify CRISPRi-validated regulatory elements. a, Overview of the CRISPRi benchmarking workflow. Variants from GWAS, eQTL, or pooled datasets were scored with ChromBPNet, aggregated to the level of candidate cisregulatory elements (cCREs), and tested for association with CRISPRi-validated target-gene status. Aggregation used the strongest predicted effect, mean predicted effect, or fraction of variants predicted to increase accessibility. Robustness was assessed across alternative filtering and analysis parameters. **b,** Three highest-performing significant models for each data source (GWAS, eQTL, and pooled). Points show odds ratios and horizontal bars show 95% confidence intervals. The dashed line marks no association (OR = 1). **c,** Distribution of odds ratios across all tested parameter combinations, grouped by aggregation method and data source. Violin widths indicate density, horizontal lines indicate medians, and thick bars indicate interquartile ranges. The dashed line marks no association. **d,** Example K562 cCRE containing a CTCF motif identified from ChromBPNet contribution scores. *Top*, contribution score across the cCRE. *Bottom*, K562 CTCF ChIP-seq signal across the same region. The shaded region marks the CTCF motif instance identified by Fi-NeMo. *Inset*, motif logo derived from ChromBPNet contribution scores. **a** created in BioRender. Kazemi, M. (2026) https://BioRender.com/u2fjd32.

cCREs with larger predicted accessibility gains were more likely to have a validated target gene in our primary analysis. However, this association was weak and sensitive to analytical choices. Across a range of alternative specifications, the estimated effect was usually positive but varied substantially in magnitude, and the number of nominally significant results was close to that expected by chance (Fig. 4b,c; Methods; Additional file 1: Fig. S6, Note S7). We therefore interpret the association as suggestive rather than evidence that predicted accessibility gain reliably identifies gene-linked cCREs.

The strongest signal came from the magnitude of predicted accessibility gain rather than the fraction of variants predicted to increase accessibility. The association was also stronger among distal cCREs, where genomic proximity provides less information for linking regulatory elements to genes. These patterns suggest that accessibility predictions may help prioritize candidate regulatory elements, particularly when other evidence is limited, although larger CRISPRi datasets will be needed to assess this more rigorously (Additional file 1: Note S7).

Together, these results show that ChromBPNet predictions provide evidence that a variant may alter chromatin accessibility, but they are much less informative for identifying the target gene affected by that variant.

### Candidate CREs containing a CTCF motif are less likely to have a CRISPRi-validated target gene

We next asked whether particular transcription factor motifs distinguish cCREs with validated target genes from those without. Among motifs identified from ChromBP- Net contribution scores (Methods), CTCF showed the strongest association. cCREs containing a confident CTCF motif were less likely to have a CRISPRi-validated target gene, and this relationship remained after accounting for other motifs, predicted accessibility, chromatin activity, and additional genomic features (Additional file 1: Fig. S7, Notes S8-S9). The association was consistent across CRISPRi studies and was supported by K562 CTCF ChIP-seq data (Fig. 4d; Methods). In contrast, a second motif group resembling NFE2/NRF2 showed a signal only when data from different studies were pooled and not in comparisons within studies. Because these sequences also matched several related bZIP motifs, we did not interpret this association further (Additional file 1: Note S8).

CTCF is best known for its role in chromatin organization rather than transcrip-tional activation [23]. Consistent with this function, CTCF motifs marked accessible elements that were less likely to be linked to a measurable target gene by CRISPRi. These results further support the idea that chromatin accessibility alone does not distinguish gene-regulatory cCREs from other classes of accessible elements.

### Predicted variant effects and trait-level chromatin enrichment across the blood lineage are released as an interactive resource

We release an interactive resource, BloodNNet (https://bloodnnet.ccbr.utoronto.ca/), that provides both variant-level and trait-level views of regulatory activity across the hematopoietic lineage (Fig. 5).

**Fig. 5.**
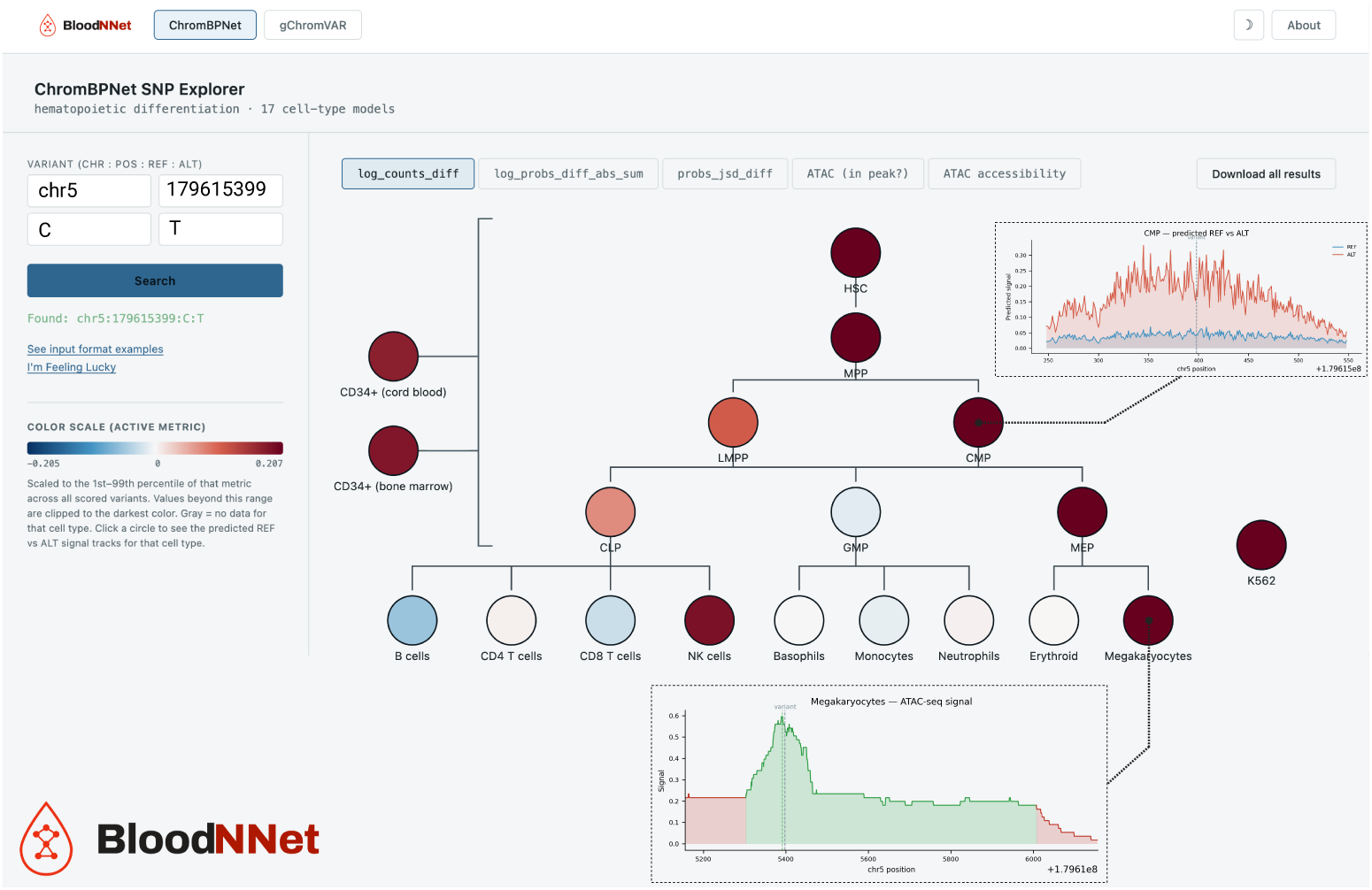
The BloodNNet web resource. Screenshot of BloodNNet (https://bloodnnet.ccbr.utoronto.ca/) showing ChromBPNet predictions for a variant across the hematopoietic differentiation hierarchy. Each node represents a blood cell type and is colored by the selected prediction metric, here the predicted log fold change in chromatin accessibility between the alternate and reference alleles (log counts diff). Red indicates increased predicted accessibility and blue indicates decreased predicted accessibility. Selecting a cell type displays the predicted accessibility profiles for the reference and alternate alleles at that locus (*top right* ). The observed ATAC-seq signal for the selected cell type can also be displayed (*bottom*), allowing predicted effects to be viewed in the context of local chromatin accessibility.

The variant page reports ChromBPNet predictions of a variant’s effect on chromatin accessibility across blood cell types. Users can query single-nucleotide variants and indels and visualize both predicted accessibility profiles and summary effect metrics across the hematopoietic differentiation hierarchy. Several million variants from CAUSALdb [24], ClinVar [25] and the eQTL Catalogue [22] have been precomputed, while additional variants can be scored on demand.

The trait page reports gChromVAR [8] enrichment of fine-mapped variants across blood cell types for 10,375 traits drawn from CAUSALdb. Trait enrichments and variant predictions are displayed on the same hematopoietic tree, allowing direct comparison between the cell types in which a variant is predicted to act and the cell types enriched for variants associated with a trait.

All predictions, enrichment scores, and metadata are available for download. The resource is intended to support prioritization of candidate regulatory variants and interpretation of blood cell trait associations in their cellular context.

## Discussion

Our results define what chromatin accessibility predictions can and cannot contribute to noncoding variant interpretation. Predicted effects are strongly structured by cell type and lineage and show agreement with statistical fine-mapping. In contrast, predicted accessibility changes provide only weak evidence about which genes candidate regulatory elements control. Together, these findings place chromatin accessibility predictions within a broader framework for variant prioritization.

The clearest result is that predicted regulatory effects follow the hematopoietic hierarchy. Variants active in more cell types tended to have larger predicted effects, replicating observations made across diverse tissues by Marderstein et al. [7]. The same pattern was observed with AVI [18], suggesting that it is not specific to ChromBPNet. One practical consequence is that genome-wide prioritization schemes preferentially select broadly acting variants and may underprioritize variants with strong effects in a single lineage. Cell-type-resolved predictions remain important for identifying these variants.

A differentiation hierarchy adds information that is not available when cell types are treated as independent contexts. Variants were preferentially shared between developmentally related cell types, and sharing declined with lineage distance. When effects were shared between progenitors and descendants, their direction was almost always preserved. This pattern is consistent with a model in which sequence variation determines the direction of effect, whereas cellular context determines its magnitude. More broadly, it suggests that cell-type-specific predictions contain information about developmental relationships that is lost when effects are summarized across tissues.

Comparison with fine-mapping helps clarify what ChromBPNet contributes. Agreement increased with fine-mapping confidence, but most variants were still nominated by only one method. The disagreeing classes differed in biologically sensible ways. Fine-mapping-only variants were enriched for coding and transcript-associated variation, whereas ChromBPNet-only variants were enriched for regulatory elements, accessible chromatin, and lineage-concordant activity. These observations suggest that ChromBPNet captures regulatory signals that are largely orthogonal to those used by association-based methods. They also suggest that chromatin accessibility predictions could provide useful priors for future fine-mapping approaches.

Accessibility predictions carried much less information about which genes a regulatory element controls. Candidate regulatory elements containing variants with larger predicted accessibility gains showed only weak enrichment for validated target genes in CRISPRi datasets. The association was directionally consistent across analyses but too small and unstable to support strong conclusions. Chromatin accessibility predictions therefore appear more informative for identifying where a variant is likely to act than for identifying the downstream gene affected by that variant.

The strongest motif-level signal involved CTCF. Candidate regulatory elements containing a CTCF motif were less likely to have a CRISPRi-validated target gene, and this association was consistent across studies and analytical approaches. CTCF is best known for its role in chromatin organization rather than transcriptional activation [23]. Consistent with this function, CTCF motifs marked accessible elements that were less likely to regulate a measurable target gene in CRISPRi screens. More generally, these results emphasize that chromatin accessibility and gene regulation are related but distinct properties.

Several limitations should be considered. Our conclusions are based primarily on a single model family and therefore concern ChromBPNet rather than sequence-based predictors in general. The hematopoietic hierarchy remains incompletely sampled, and neutrophil and basophil models did not pass quality control. The CRISPRi analyses were limited to K562 cells and to a relatively small number of validated regulatory elements. Finally, the lineage concordance of model-only variants supports a regulatory interpretation but does not establish causality.

Together, these results show that chromatin accessibility predictions are most informative for identifying the cell types in which a noncoding variant is likely to act, less informative for determining which gene it affects, and most powerful when combined with complementary sources of evidence. We release these predictions through the BloodNNet resource so that investigators can compare variant-level regulatory predictions with trait-level genetic enrichments across the hematopoietic hierarchy. Experimental characterization of the high-confidence variants identified here, together with extension of the framework to additional cell types and gene-linking strategies, are natural next steps.

## Conclusions

Chromatin accessibility predictions are most informative for identifying the cell types in which a noncoding variant acts, less informative for identifying its target gene, and most powerful when combined with complementary sources of genetic evidence.

## Methods

### Fine-mapping and background variant sets

Posterior inclusion probabilities (PIP) and 95% credible sets were taken from the published fine-mapping of 29 blood cell indices in UK Biobank by Vuckovic et al. [2], which used FINEMAP v1.3.1 [26]. Coordinates were lifted from GRCh37 to GRCh38, and variants were retained at PIP *≥* 0.001. The 29 traits comprise 14 red cell indices (hematocrit, hemoglobin, MCH, MCHC, MCV, MSCV, RBC count, RDW, reticulocyte count and percentage, immature reticulocyte fraction, high light-scatter reticulocyte count and percentage, mean reticulocyte volume), 4 platelet indices (platelet count, plateletcrit, mean platelet volume, platelet distribution width) and 11 white cell indices (white cell count, and neutrophil, lymphocyte, monocyte, eosinophil and basophil counts and percentages). When a variant appeared more than once for a trait (separate credible sets) the maximum posterior was kept. This gave 400,192 variant-trait pairs covering 194,199 unique variants.

HapMap 3 [27] was used as a trait-agnostic background set. Coordinates were lifted from NCBI build 36 to GRCh38. Indels, and variants too close to a chromosome end to allow the 2,114 bp ChromBPNet input window, were removed. For multiallelic sites, reference and alternate alleles were matched to the Vuckovic et al. [2] summary statistics so that model predictions and association statistics referred to the same allele. No allele-frequency filter was applied. The final background set contained 1,404,991 variants.

Matching the scored variants against the Vuckovic summary statistics showed that 99.98% had an association result for at least one trait. The union of the fine-mapped and HapMap 3 datasets contained 1,565,538 distinct variants, including 36,194 present in both.

### ChromBPNet models, variant scoring and activity calls

ChromBPNet models were trained for 17 blood cell types from public ATAC-seq data (Additional file 2: Table S3; Fig. 1a), including hematopoietic stem cells, multipotent progenitors, lymphoid-primed multipotent progenitors, common myeloid progenitors, granulocyte–monocyte progenitors, megakaryocyte–erythroid progenitors, common lymphoid progenitors, CD34^+^ cells from bone marrow and from umbilical cord blood, erythroid cells, megakaryocytes, monocytes, B cells, CD4^+^ T cells, CD8^+^ T cells, NK cells and K562. Primary hematopoietic populations came from GSE74912 [28], megakaryocytes from GSE96772 [29] and K562 from GSE213295 [30]. Raw FASTQ files were downloaded from GEO and processed with nf-core/atacseq (v2.1.2) [31] under Nextflow v25.10.3 [32]. Biological and technical replicates were merged into a single alignment and peak set for each cell type. One ChromBPNet bias model and one bias-factorized accessibility model were then trained per cell type following the standard ChromBPNet workflow [5] with five-fold cross-validation. Models were also trained for basophils (GSE138610 [33]) and neutrophils (GSE153520 [34]), but neither passed model quality control and both were excluded from all analyses. These models remain available through BloodNNet for exploratory use.

Variants were scored in every cell-type model using the ChromBPNet variantscoring workflow (kundajelab/variant-scorer [5]). For each allele, the model predicts base-resolution chromatin accessibility across a 1 kb window centered on the variant. The workflow reports three effect metrics: log counts diff, log probs diff abs sum, and probs jsd diff. Unless otherwise noted, analyses used the predicted log_2_ fold change in accessibility (logFC). Magnitude analyses used *|*logFC*|*, whereas analyses of direction used signed logFC. Jensen-Shannon divergence (JSD) was used only in threshold-sensitivity analyses.

Statistical significance was assessed using an empirical null generated from dinucleotide-preserving shuffles of the 2,114 bp sequence surrounding each variant, with the reference or alternate allele inserted at the center. Observed and null scores were averaged across the five folds before testing. For each cell type the one-sided empirical P was calculated as (1 + #*{*null *≥* observed*}*)*/*(1 + *n*_null_) using a null distribution of 100,000 fold-averaged shuffled sequences. The smallest attainable P value was therefore 1/100,001 *≈* 1.0 *×* 10*^−^*^5^.

The scored dataset comprised 1,565,538 distinct variants from the union of the finemapped and HapMap 3 variant sets. A variant was considered tested if it overlapped the consensus accessible-region set (302,830 variants) and accessible in a given cell type if it overlapped that cell type’s peak set. Analyses of effect magnitude were restricted to the 242,307 variants accessible in at least one cell type.

A variant was considered active in a cell type if it was accessible in that cell type and had an empirical P *<* 0.01. Variants active in at least one cell type were classified as active overall. Cross-cell-type comparisons were normalized by the number of accessible variants in each cell type. Sensitivity analyses repeated all downstream analyses using nine alternative calling rules: BH-FDR *<* 0.1, 0.05, and 0.01 based on logFC; empirical P *<* 0.05, 0.01, and 0.001 based on logFC; and empirical P *<* 0.05, 0.01, and 0.001 based on logFC *×* JSD. BH-FDR was calculated separately within each cell type using the variants accessible in that cell type.

### Breadth and regulatory magnitude

Breadth was defined as the number of cell types (of 17) in which a variant was both accessible and active. Following Marderstein et al. [7], variants were grouped as inactive, active in 1 cell type, active in 2–13 cell types, or active in at least 14 cell types (*≥*80% of cell types). Regulatory magnitude was defined as the maximum absolute predicted effect (*|*logFC*|*) across all cell types in which a variant was accessible, regardless of whether it was called active. Group comparisons report mean magnitude with 95% confidence intervals, together with ordinary least squares (OLS) contrasts against the inactive group and Welch tests between active groups. We also divided variants into 10 equal-count bins of regulatory magnitude and compared mean breadth across bins.

To evaluate the relationship between breadth and effect size, we calculated four Spearman correlations restricted to variants with breadth *>*0: (i) maximum effect size across active cell types versus breadth; (ii) maximum effect size across all cell types versus breadth; (iii) a leave-one-out analysis comparing effect size in cell type c with breadth calculated after excluding cell type c; and (iv) effect size in cell type c versus the number of other cell types in which the variant was accessible. The leave-one-out analysis was the primary estimator because it separates the effect used as the outcome from the cell-type activity contributing to breadth.

To quantify the amount of breadth-magnitude association expected from the definition of breadth alone, we generated 10,000 permutations by shuffling effect sizes and significance calls jointly among accessible variants within each cell type. Breadth and all summary statistics were recalculated for each permutation, and results are reported relative to the null mean. Permutation P values were calculated using a one-sided test with a +1 correction, giving a minimum attainable P value of 1/10,001.

All analyses were repeated at five activity thresholds (P *<* 0.05, 0.01, 0.001, 10*^−^*^4^, and 10*^−^*^5^). Confidence intervals for the leave-one-out analysis were calculated using the number of distinct variants within each breadth stratum because a single variant can contribute observations from multiple cell types.

### Comparison with the AlphaGenome Variant Impact score

AlphaGenome Variant Impact (AVI) scores were downloaded from the AlphaGenome Atlas [18] (https://deepmind.google.com/science/alphagenome/downloads) and queried using tabix (htslib 1.21 [35]) at the positions of the 242,307 variants accessible in at least one cell type. Variants were matched by chromosome, position, and alleles, yielding 241,575 exact matches (99.7%). Analyses used the PHRED-scaled AVI score, for which values of 10, 20, and 30 correspond to the top 10%, 1%, and 0.1% of scored variants genome-wide [18]. Results were unchanged when using raw AVI scores.

Breadth classes and regulatory magnitude were defined from ChromBPNet predictions as described above. Comparisons among breadth classes used the same framework as the breadth analyses, including group means, confidence intervals, OLS contrasts, and pairwise tests. We additionally calculated the fraction of variants exceeding PHRED thresholds of 10, 20, and 30, odds ratios relative to inactive variants, and AUROC values for class discrimination.

Because broader variants tend to have larger predicted effects and occur in more accessible chromatin, all primary comparisons were repeated after matching variants for regulatory magnitude and accessibility breadth. The primary analysis compared variants within strata defined by these two variables. Additional adjustment approaches are described in Additional file 1: Note S2.

To assess the effect of AlphaGenome cell-type coverage, each hematopoietic cell type was classified as directly profiled, represented by bulk CD34+ data, or not represented by a closely matching accessibility dataset. AVI contrasts were then repeated within and between these coverage groups, including analyses restricted to variants active in profiled or non-profiled cell types.

Sensitivity analyses repeated all comparisons across alternative activity thresholds, included allele-swapped AVI matches, excluded coding and canonical splice-site variants defined by VEP [36], and stratified variants by distance to the nearest proteincoding transcription start site (GENCODE v47 [37]). Additional details are provided in Additional file 1: Note S2.

### Annotation with ENCODE candidate cis-regulatory elements

Variants were annotated using version 4 of the ENCODE candidate cis-regulatory element (cCRE) registry [38] for GRCh38 (2,348,854 elements). These comprised distal enhancer-like (dELS), proximal enhancer-like (pELS), promoter-like (PLS), chromatin-accessible, CTCF-associated, transcription-factor-associated, and related subclasses. A variant was assigned the class of the cCRE containing it or classified as “no cCRE” if no overlap was present. Where multiple overlaps occurred, classes were prioritized in the order PLS *>* pELS *>* dELS *>* CA-H3K4me3 *>* CA-CTCF *>* CA- TF *>* CA *>* TF. In practice, no variant overlapped more than one cCRE. Overall, 51.7% of the 242,307 accessible variants overlapped a cCRE.

Associations between breadth categories and cCRE classes were evaluated using Fisher’s exact tests applied to 2 *×* 2 contingency tables. Two comparisons were performed for each breadth category and cCRE class: against inactive variants and against the remaining active variants pooled. Odds ratios and 95% confidence intervals were calculated from the log odds ratio standard error, and Benjamini-Hochberg correction was applied across all tests. Classes represented by fewer than five variants in the category of interest were not tested.

To assess whether cCRE composition differed among the three active breadth categories, we applied a chi-square test of independence to the category-by-cCRE contingency table. Because several expected cell counts were below five, statistical significance was additionally assessed by permutation. cCRE labels were permuted across active variants 2,000 times while preserving category and class totals, and the empirical P value was calculated using a +1 correction.

### Lineage analyses

The 14 primary hematopoietic populations were represented as a differentiation graph spanning stem cells, progenitors, and mature blood cell lineages (Fig. 2a). Distances between cell types were defined as shortest-path distances on the undirected graph. K562 and the two bulk CD34^+^ populations were excluded from lineage analyses.

For variants active in at least two cell types, we quantified lineage coherence using two metrics: (i) the fraction of variants whose active cell types formed a connected subgraph and (ii) the mean pairwise lineage distance among active cell types. Both statistics were compared with a curveball null model preserving each variant’s breadth and each cell type’s number of active variants. Significance was assessed using 1,000 permutations and one-sided empirical P values.

To quantify sharing between cell types, we counted variants active in both members of each of the 91 cell-type pairs and compared those counts with expectations under independence among variants accessible in both cell types. We then tested whether the excess sharing declined with lineage distance. Statistical significance was assessed by permuting cell-type positions on the lineage graph 5,000 times.

To determine whether lineage effects could be explained by overall similarity between cell types, the pairwise model was repeated with an additional covariate representing the correlation of absolute predicted logFC values between cell types, calculated using variants accessible in both cell types but active in neither. Additional robustness analyses are described in Additional file 1: Note S4.

For each parent-child edge in the lineage graph, variants active in both cell types were compared for consistency in the sign of their predicted effects. The proportion of variants with opposing effect directions was reported with Jeffreys 95% confidence intervals.

### Comparison with statistical fine-mapping

Variants were classified as fine-mapping-nominated if they had PIP *≥* 0.5 for at least one of the 29 blood cell traits and as model-nominated if they were active in at least one of the 17 cell types. All variants with both a model prediction and fine-mapping result were assigned to one of four classes: both, fine-mapping-only, model-only, or neither.

Associations between fine-mapping and model nominations were evaluated using Fisher’s exact tests and odds ratios. The primary analysis was restricted to the 36,200 variants accessible in at least one cell type. We additionally repeated the analysis after comparing variants with similar accessibility breadth. Additional analyses examining fine-mapping coverage, alternative universes, and accessibility stratification are described in Additional file 1: Note S5.

To assess graded agreement, variants were grouped by their maximum PIP across the 29 traits, and the proportion nominated by ChromBPNet was calculated within each bin. Trends across ordered PIP bins were evaluated using a linear-by-linear association test.

We compared the empirical-P activity definition with two alternative nomination rules: a fixed effect-size threshold (*|*logFC*| ≥* 0.5) and a magnitude-only rule selecting the top-ranked variants by *|*logFC*|*. Rule concordance and enrichment analyses are described in Additional file 1: Note S5.

High-confidence variants were defined as variants with PIP *≥* 0.5 and empirical P *<* 0.001 in at least one accessible cell type. Nearest genes were assigned by genomic distance, and regulatory breadth was defined as the number of cell types in which the variant was active.

### Consequence annotation, active-allele quantile and cell-type concordance

Variants were annotated with Ensembl VEP v115.2 [36] using GRCh38.p14, GEN- CODE v47 [37], dbSNP 156 [39], the Ensembl regulatory build 1.0 [40], and gnomAD v4.1 [41]. Regulatory annotations were enabled using the --regulatory option. Variants carrying multiple annotations were counted in each applicable consequence class.

Consequence enrichments were evaluated for fine-mapping-only, model-only, and doubly nominated variants relative to either all non-nominated variants or the subset accessible in at least one cell type. Odds ratios and confidence intervals were calculated from 2 *×* 2 contingency tables, and P values were adjusted using the Benjamini- Hochberg procedure. Additional details and sensitivity analyses are provided in Additional file 1: Note S6.

The active-allele quantile (AAQ) was defined as the genome-wide quantile of the allele predicted to have higher accessibility [5]. For each variant, AAQ was taken from the cell type with the largest absolute predicted effect. Differences among nomination classes were assessed using pairwise rank-based comparisons, with false-discovery-rate correction across all tests.

For cell-type concordance analyses, a variant was linked to a trait if it appeared in a PIP *≥* 0.001 credible set for that trait. Trait lineages were defined as described in Additional file 1: Note S6. A variant-trait pair was considered concordant if the variant was active in at least one cell type belonging to the relevant lineage. Concordance was evaluated against a permutation null that preserved both variant activity patterns and the number of trait assignments per variant while randomizing the variant-trait correspondence. Significance was assessed using 10,000 permutations. Additional lineage definitions, concordance analyses, and sensitivity analyses are provided in Additional file 1: Note S6.

### CRISPRi benchmark in K562 cells

Candidate cis-regulatory elements (cCREs) were compiled from published K562 CRISPRi studies [9–17] and combined into a single set of hg38 intervals with a binary indicator of whether each element had at least one experimentally validated target gene. Because the studies differed in design, significance was defined using the primary criterion reported by each study. The integrated ENCODE resource [16] was used to obtain element-level calls from FlowFISH (Nasser et al. [9]), Perturb-seq data (Gasperini et al. [10]), and TAP-seq data (Schraivogel et al. [15]) datasets. Positive elements were those marked as significant. For the Gasperini screen, we used the independent SCEPTRE [42] reanalysis of Ghatan et al. [17] rather than the reanalysis included in the integrated resource. From the ENCODE4 meta-analysis of bulk CRISPRi screens (Yao et al. [14]), we retained K562 elements and used the reported target-gene indicator. From ter Weele et al. [13], we included the Perturb-seq followup dataset and considered a site-gene pair positive when the reported false-discovery rate was below 0.05. The genome-wide fitness screen component was not used. Reilly et al. [11] assayed a limited set of loci and genes using HCR-FlowFISH. Because the study was not designed to identify all possible targets, we retained only the reported positive elements and did not treat unvalidated elements as negatives. For our STING-seq study (Morris et al. [12]), guide RNAs were aligned to hg38 and converted to regulatory-element intervals by extending 1 kb on either side of the aligned position. Elements were considered positive if any reported association passed a false-discovery-rate threshold of 0.05.

The final dataset contained 9,205 cCREs, of which 1,259 had at least one validated target gene (Additional file 2: Table S4). Each element was annotated with distance to the nearest protein-coding transcription start site (TSS), overlap with K562 ATAC-seq peaks, and GC content.

Two variant sets were used for ChromBPNet K562 predictions. The first contained fine-mapped blood-cell-trait variants from Vuckovic et al. [2]. The second contained fine-mapped (PIP *≥* 0.05) eQTL variants from 39 blood and immune datasets in the eQTL Catalogue [22] (Additional file 2: Table S5). After combining and deduplicating the two datasets, the pooled set contained 1,447,084 unique variants.

For each cCRE, variants falling within the element boundaries were identified and summarized using three scores: the strongest predicted accessibility effect, the mean predicted effect, and the fraction of variants predicted to increase accessibility. Elements containing no variants were excluded from a given analysis. We then tested whether these element-level scores were associated with the presence of a validated target gene while adjusting for distance to the nearest protein-coding TSS. The primary analysis used the pooled variant set, no effect size threshold, no accessibility filter, and elements located more than 10 kb from the nearest TSS. The strongest predicted effect was used as the primary summary metric. To assess robustness, the analysis was repeated across alternative variant sets, aggregation rules, effect size thresholds, accessibility filters, and TSS-distance thresholds. Details of the specification sweep and power calculations are provided in Additional file 1: Note S7.

### Motif analysis of CRISPRi cCREs

Base-resolution contribution scores were computed for each K562 cCRE using the K562 ChromBPNet model and DeepSHAP [43, 44], following the contribution-scoring workflow implemented in kundajelab/variant-scorer [5]. Scores were calculated across a 2,114-bp window centered on the midpoint of each element.

Motif instances were identified using Fi-NeMo [45] and a pre-existing K562 ChromBPNet TF-MoDISco-lite motif compendium [46] generated independently of the CRISPRi dataset. Each motif instance was assigned a putative transcription-factor identity using the top TomTom match [47]. For the primary analysis, we retained motif instances that fell entirely within the annotated cCRE boundaries and had a confident motif match. A secondary analysis used all motif instances weighted by motif-match confidence.

For each cCRE and transcription-factor label, we quantified motif activity using motif presence, motif count, and the summed contribution score across motif instances. Additional representations based on element-length normalization and confidence weighting were evaluated in sensitivity analyses (Additional file 1: Note S8).

We first compared motif frequencies between cCREs with and without validated target genes. We then fit multivariable models relating target-gene status to motif features while adjusting for ABC score [48], chromatin accessibility, GC content, distance to the nearest transcription start site, sequence constraint, element length, and predicted accessibility. Additional details and sensitivity analyses are described in Additional file 1: Notes S8 and S9.

To assess robustness, motif analyses were repeated after excluding the Reilly dataset and were additionally evaluated within individual studies. Study-specific analyses and validation of CTCF motif calls using K562 CTCF ChIP-seq data are described in Additional file 1: Notes S8 and S9.

### Validation against K562 CTCF ChIP-seq

We used ENCODE IDR-thresholded K562 CTCF ChIP-seq peaks [49] (experiment ENCSR000DWE, file ENCFF736NYC; GRCh38) to validate CTCF motif instances identified by Fi-NeMo. Peak summits were taken from the narrowPeak summit annotations. Fold-enrichment tracks from ENCFF024JKM were used for visualization in Fig. 4d.

CTCF and NFE2/NRF2 motif instances were taken directly from the Fi-NeMo analyses described above. For the primary analysis, only confident motif instances located entirely within the annotated cCRE boundaries were considered. A motif instance was considered supported by ChIP-seq if its center fell within a CTCF peak. Enrichment over expectation was assessed by randomly repositioning motif instances within their own cCREs while preserving motif width. Additional analyses compared distances to peak summits and overlap frequencies at both the motif and cCRE levels. To evaluate the relationship between CTCF occupancy and regulatory activity, we compared motif frequencies between cCREs with and without CRISPRi-validated target genes. We additionally evaluated CTCF motifs with and without ChIP-seq support, stratified analyses by distance to the nearest protein-coding transcription start site, and repeated comparisons within individual studies. Details of these analyses are provided in Additional file 1: Note S9.

Finally, we examined motif orientation and its relationship to target-gene status by comparing motif-strand distributions, cCRE orientation classes, and motif orientation relative to nearby ChIP-seq summits.

## Supplementary information

**Additional file 1:** Supplementary Notes S1–S9, Tables N1.1–N8.3 and Supplementary Figures S1–S7.

**Additional file 2:** Supplementary Tables S1–S10.

**Table S1.** High-confidence variants supported by both ChromBPNet and finemapping

**Table S2.** All model-nominated (active) variants

**Table S3.** ATAC-seq samples used to train ChromBPNet models

**Table S4.** Curated K562 CRISPRi candidate regulatory elements

**Table S5.** eQTL Catalogue datasets used for the CRISPRi analysis

**Table S6.** Consequence enrichment of nominated variants (full results)

**Table S7.** Motif association with CRISPRi-validated target genes (30 motif labels *×* 5 schemes)

**Table S8.** Multivariable model of target-gene status: motif presence scheme

**Table S9.** Multivariable model of target-gene status: motif importance scheme

**Table S10.** CTCF and NFE2/NRF2 motif frequency by CRISPRi study

## Declarations

### Ethics approval and consent to participate

Not applicable.

### Consent for publication

Not applicable.

### Availability of data and materials

ChromBPNet models for 19 hematopoietic cell types are available on Zenodo at https://doi.org/10.5281/zenodo.24091438. Custom scripts for analyzing data and generating figures are available on GitHub at https://github.com/jmorrislab/BloodNNet.

### Authors’ contributions

J.A.M. conceived the study and supervised the research. M.K. performed all computational analyses, generated the figures and tables, and developed the BloodNNet resource. J.A.M. and M.K. interpreted the results and wrote the manuscript.

## Acknowledgements

We thank the entire Morris laboratory for support and advice. We thank Dr. Michael Wainberg and Dr. Selin Jessa for helpful discussions in interpreting noncoding variant effects.

