## Additional file 1 for "Chromatin accessibility predictions reveal lineage-structured effects of noncoding variants across hematopoiesis"

#### Additional file 1: Supplementary Notes, Tables and Figures

###### Tables in the Supplementary Notes

#### Supplementary Note S1: Robustness of the breadth–magnitude relationship

**S1.1 Distribution of activity.** Of the 242,307 variants accessible in at least one cell type, 13,608 were active in at least one cell type (Fig. S1a,b). Most active variants were restricted to a single cell type (7,743; 56.9%), while only 77 variants (0.57%) were active in at least 14 of the 17 cell types.

**S1.2 Sensitivity to activity threshold.** To assess whether the breadth-magnitude relationship was driven by false-positive activity calls, we repeated the analysis across progressively stricter activity thresholds. The leave-one-out correlation remained consistently positive across thresholds (Table N1.1; Fig. S1c), arguing against the relationship being driven by false-positive calls.

**S1.3 Alternative estimators of the breadth-magnitude relationship.** Because breadth is partly defined using the same information used to measure effect size, estimators that share a cell type between predictor and outcome can produce inflated correlations. We therefore used the leave-one-out estimator as the primary analysis. The leave-one-out correlation remained strongly positive and greatly exceeded its permutation null, whereas breadth defined by chromatin accessibility alone showed no positive association with effect size (Table N1.2).

**S1.4 Magnitude by breadth group.** Regulatory magnitude increased with breadth, from a mean of 0.442 among single-cell-type variants to 1.163 among variants active in at least 14 cell types (Table N1.3).

#### Supplementary Note S2: Breadth and the AlphaGenome Variant Impact score

**S2.1 AlphaGenome Variant Impact scores.** The AlphaGenome Variant Impact (AVI) score is a genome-wide prioritization score derived from AlphaGenome predictions together with coding and conservation features [18]. We used AVI as an independent benchmark to test whether the breadth gradient observed with ChromBPNet was also present in a model trained for a different objective. Of the 242,307 variants accessible in at least one cell type, 241,575 (99.7%) matched AVI exactly by position and alleles. Match rates were similar across breadth classes.

**S2.2 AVI increases with regulatory breadth.** AVI increased with breadth in the same direction as ChromBPNet regulatory magnitude (Table N2.1). Variants active in more cell types had higher mean AVI scores, with the highest scores observed among variants active in at least 14 cell types.

**S2.3 Genome-wide AVI cutoffs preferentially select broadly active variants.** Broadly active variants were more likely to exceed genome-wide AVI thresholds (Table N2.2). Although single-cell-type variants comprised 56.9% of active variants, they represented only 27.6% of active variants exceeding PHRED $\geq$ 20 (top 1% genome-wide).

**S2.4 Matching for regulatory magnitude and accessibility breadth.** Because broader variants tend to have larger predicted effects and occur in more accessible chromatin, we repeated the AVI comparisons after matching variants for both regulatory magnitude and accessibility breadth. The difference between

**Table N1.1** Breadth–magnitude statistics across calling thresholds.

| **Threshold (empirical P)** | **Active variants** | **Pairs** | **At breadth 1** | **$\rho$ panel** | **$\rho$ leave-one-out** | **$\rho$ accessibility** |
|---|---|---|---|---|---|---|
| $< 0.05$ | 45,367 | 120,010 | 53.4% | +0.535 | +0.325 | −0.040 |
| $< 0.01$ (primary) | 13,608 | 32,519 | 56.9% | +0.581 | +0.386 | −0.024 |
| $< 0.001$ | 2,545 | 5,721 | 58.4% | +0.610 | +0.384 | −0.001 |
| $< 10^{-4}$ | 461 | 852 | 65.9% | +0.606 | +0.450 | +0.052 |
| $< 10^{-5}$ | 88 | 118 | 80.7% | +0.519 | +0.412 | −0.049 |

**Table N1.2** Estimators of the breadth–magnitude correlation against the permutation null.

| Estimator | Observed $\rho$ | Null mean (SD) | Excess | Permutation P |
|---|---|---|---|---|
| Full panel | +0.581 | 0.221 (0.005) | +0.360 | $< 10^{-4}$ |
| Leave-one-out | +0.386 | 0.029 (0.005) | +0.358 | $< 10^{-4}$ |
| Accessibility breadth | −0.024 | 0.077 (0.006) | −0.101 | 1.0 |

**Table N1.3** Regulatory magnitude by breadth group.

| Group | n | Mean magnitude [95% CI] | Difference vs inactive [95% CI] |
|---|---|---|---|
| Inactive | 228,699 | 0.055 [0.0546, 0.0551] | – |
| One cell type | 7,743 | 0.442 [0.437, 0.448] | +0.387 [0.382, 0.393] |
| Two to thirteen | 5,788 | 0.739 [0.729, 0.749] | +0.684 [0.674, 0.694] |
| $\geq$ 14 | 77 | 1.163 [1.055, 1.272] | +1.109 [1.00, 1.22] |

**Table N2.1** AlphaGenome Variant Impact (AVI) PHRED score by breadth group.

| Group | n | Mean AVI PHRED [95% CI] | Median | Difference vs inactive [95% CI] |
|---|---|---|---|---|
| Inactive | 228,009 | 4.53 [4.51, 4.55] | 3.06 | – |
| One cell type | 7,713 | 9.20 [9.08, 9.33] | 9.07 | +4.67 [4.57, 4.78] |
| Two to thirteen | 5,776 | 12.73 [12.59, 12.88] | 12.92 | +8.20 [8.08, 8.32] |
| $\geq$ 14 | 77 | 18.28 [17.10, 19.45] | 19.09 | +13.75 [12.71, 14.78] |

**Table N2.2** Fraction of each breadth group passing genome-wide AVI cutoffs.

| Cutoff | Inactive | 1 cell type | 2–13 cell types | $\geq$ 14 cell types | 1 cell type share of active passing |
|---|---|---|---|---|---|
| PHRED $\geq$ 10 (top 10%) | 12.5% [12.4, 12.6] | 43.2% [42.1, 44.3] | 69.5% [68.3, 70.7] | 93.5% [85.7, 97.2] | 44.9% |
| PHRED $\geq$ 20 (top 1%) | 0.7% [0.7, 0.8] | 2.8% [2.5, 3.2] | 9.3% [8.5, 10.0] | 42.9% [32.4, 54.0] | 27.6% |

broader variants and single-cell-type variants persisted after matching (Table N2.3; Fig. S1e), indicating that the breadth gradient is not explained solely by differences in ChromBPNet effect size or accessibility breadth.

**S2.5 AlphaGenome coverage.** To determine whether the residual breadth effect could be explained by differences in AlphaGenome cell-type coverage, we repeated the matched comparison separately among variants active in at least one directly profiled

**Table N2.3** AVI differences between breadth groups at matched magnitude and accessibility.

| Contrast | Matched on | Difference [95% CI] | Variants used |
|---|---|---|---|
| One vs inactive | magnitude | −0.21 [−0.43, 0.00] | 4,410 vs 17,911 |
| One vs inactive | magnitude × accessibility | −0.14 [−0.35, 0.08] | 4,384 vs 16,711 |
| Two to thirteen vs one | magnitude | +1.48 [1.26, 1.69] | 5,771 vs 7,455 |
| Two to thirteen vs one | magnitude × accessibility | +1.16 [0.92, 1.40] | 5,156 vs 6,923 |
| Two to thirteen vs one | spline + accessibility | +1.07 [0.81, 1.32] | all active |
| ≥ 14 vs two to thirteen | magnitude × accessibility | +2.38 [1.10, 3.66] | 59 vs 213 |
| ≥ 14 vs two to thirteen | spline + accessibility | +3.08 [1.92, 4.24] | all active |
| ≥ 14 vs one | spline + accessibility | +4.15 [2.96, 5.34] | all active |

cell type and among variants active in none. The difference between broader and single-cell-type variants was similar in both groups (+1.04 [0.71, 1.37] and +1.10 [0.74, 1.46]; Fig. S1f), indicating that coverage does not explain the breadth effect on its own.

**S2.6 Sensitivity to activity threshold.** To assess whether the breadth effect was driven by false-positive activity calls, we repeated the analysis using progressively stricter activity thresholds. The difference between broader and single-cell-type variants persisted across thresholds (Table N2.4), arguing against false-positive calls as the primary explanation.

**S2.7 Additional sensitivity analyses.** Results were unchanged when including allele-swapped AVI matches, restricting analyses to noncoding variants, or stratifying variants by distance to the nearest protein-coding transcription start site:

- including swapped matches: +1.17 [0.93, 1.41]
- noncoding variants only (removing 1,014): +1.15 [0.91, 1.39]
- more than 1 kb from a protein-coding TSS: +1.12 [0.88, 1.37] (5,399 vs 7,307)
- within 1 kb: +1.78 [0.35, 3.22] (377 vs 406)

Unadjusted class means are unchanged to two decimals in the first two analyses.

#### Supplementary Note S3: Regulatory element class and breadth of activity

**S3.1 cCRE composition across breadth classes.** Variants were annotated using ENCODE cCREs (Methods). Overall, 51.7% of accessible variants overlapped a cCRE. The composition of breadth classes is shown in Table N3.1 and Fig. S2a. Moving from inactive to specific to multiple variants, the proportion of variants outside any cCRE decreased from 50.2% to 21.3% to 10.4%, while the proportion in distal enhancer-like elements increased from 33.9% to 57.3% to 66.2%. Broadly active variants (≥14 cell types) were distinguished by a high proportion of promoter-like elements (29.9%).

**S3.2 Active versus inactive variants.** Relative to inactive variants, all active categories were enriched for annotated regulatory elements and depleted outside the cCRE registry (Table N3.2; Fig. S2b). Because active variants must occur in accessible chromatin with detectable predicted effects, this comparison primarily establishes that active variants are concentrated in annotated regulatory elements.

**Table N2.4** Breadth contrasts in AVI across calling thresholds.

| **Threshold** | **One / two to thirteen / ≥ 14** | **Chance bound at breadth one** | **Mean PHRED, one vs two to thirteen** | **Matched difference, one vs inactive** | **Matched difference, two to thirteen vs one** | **Pass PHRED ≥ 20, one vs two to thirteen** | **1 cell type share, active → passing** |
|---|---|---|---|---|---|---|---|
| $P < 0.01$ | 7,713 / 5,776 / 77 | ≤ 99% | 9.20 vs 12.73 | −0.14 [−0.35, 0.08] | +1.16 [0.92, 1.40] | 2.8% vs 9.3% | 57% → 28% |
| $P < 0.005$ | 4,559 / 3,438 / 38 | ≤ 87% | 10.50 vs 14.13 | 0.00 [−0.28, 0.28] | +1.31 [1.00, 1.62] | 4.1% vs 13.0% | 57% → 29% |
| $P < 0.001$ | 1,481 / 1,049 / 7 | ≤ 55% | 13.28 vs 16.85 | +0.03 [−0.47, 0.53] | +1.91 [1.36, 2.45] | 10.2% vs 24.1% | 58% → 37% |
| FDR < 0.10 | 820 / 274 / 0 | – | 14.50 vs 18.43 | −0.40 [−0.88, 0.07] | +1.76 [0.81, 2.71] | 14.6% vs 36.1% | 75% → 55% |
| FDR < 0.05 | 198 / 25 / 0 | – | 16.83 vs 21.72 | +0.14 [−0.75, 1.03] | not stable (39 matched) | 27.8% vs 68.0% | 89% → 76% |

**Table N3.1** cCRE class of the variants in each breadth category. Cells show the percentage of the category (number of variants).

| Category | n | PLS | pELS | dELS | CA-H3K4me3 | CA-CTCF | CA-TF | CA | TF | no cCRE |
|---|---|---|---|---|---|---|---|---|---|---|
| Inactive | 228,699 | 2.7 (6,170) | 7.0 (15,911) | 33.9 (77,620) | 1.0 (2,394) | 1.7 (3,801) | 0.5 (1,162) | 2.3 (5,302) | 0.7 (1,643) | 50.2 (114,696) |
| Specific | 7,743 | 4.4 (341) | 7.8 (606) | 57.3 (4,437) | 1.5 (114) | 2.5 (191) | 1.1 (84) | 3.6 (280) | 0.6 (44) | 21.3 (1,646) |
| Multiple | 5,788 | 6.4 (371) | 10.1 (587) | 66.2 (3,834) | 1.5 (85) | 2.0 (117) | 0.9 (50) | 2.3 (134) | 0.2 (9) | 10.4 (601) |
| Shared | 77 | 29.9 (23) | 14.3 (11) | 51.9 (40) | 2.6 (2) | 1.3 (1) | 0.0 (0) | 0.0 (0) | 0.0 (0) | 0.0 (0) |

**Table N3.2** Each active category against the inactive variants: odds ratio [95% CI]; BH-adjusted q, fewer than five variants in the category not tested.

| cCRE class | Specific | Multiple | Shared |
|---|---|---|---|
| PLS | 1.66 [1.49, 1.86]; q = $2.5\times10^{-16}$ | 2.47 [2.22, 2.75]; q = $1.2\times10^{-47}$ | 15.36 [9.42, 25.04]; q = $1.9\times10^{-17}$ |
| pELS | 1.14 [1.04, 1.24]; q = 0.0052 | 1.51 [1.38, 1.65]; q = $4.6\times10^{-18}$ | 2.23 [1.18, 4.22]; q = 0.029 |
| dELS | 2.61 [2.50, 2.73]; q < $10^{-300}$ | 3.82 [3.61, 4.04]; q < $10^{-300}$ | 2.10 [1.35, 3.29]; q = 0.0023 |
| CA-H3K4me3 | 1.41 [1.17, 1.71]; q = $8.9\times10^{-4}$ | 1.41 [1.13, 1.75]; q = 0.0048 | – |
| CA-CTCF | 1.50 [1.29, 1.73]; q = $8.6\times10^{-7}$ | 1.22 [1.01, 1.47]; q = 0.0495 | – |
| CA-TF | 2.15 [1.72, 2.68]; q = $3.2\times10^{-9}$ | 1.71 [1.28, 2.27]; q = $8.8\times10^{-4}$ | – |
| CA | 1.58 [1.40, 1.79]; q = $1.5\times10^{-11}$ | 1.00 [0.84, 1.19]; q = 1.0 | – |
| TF | 0.79 [0.58, 1.07]; q = 0.15 | 0.22 [0.11, 0.41]; q = $6.4\times10^{-9}$ | – |
| No cCRE | 0.27 [0.25, 0.28]; q < $10^{-300}$ | 0.12 [0.11, 0.13]; q < $10^{-300}$ | – |

**S3.3 Comparisons among active breadth categories.** Comparisons restricted to active variants revealed clear differences among breadth classes (Table N3.3; Fig. S2c). Single-cell-type variants were enriched outside annotated cCREs and depleted in promoter-like, proximal enhancer-like, and distal enhancer-like elements. In contrast, variants active in two to thirteen cell types were enriched in distal enhancer-like elements and depleted outside the registry. The broadest variants were strongly enriched in promoter-like elements. The composition of the three active categories differed overall ($\chi^2$ = 469.4, permutation P < 0.001).

#### Supplementary Note S4: Robustness of the lineage structure

**S4.1 Adjustment for chromatin similarity.** Background similarity between cell types declined with lineage distance, from 0.701 at distance 1 to 0.460 at distance 6 (Fig. S3c). When the pairwise sharing model was adjusted for background similarity, the effect of lineage distance was substantially reduced and reversed in sign (Table N4.1), consistent with chromatin similarity accounting for much of the observed distance effect.

**S4.2 Permutation-based inference.** The distance effect remained outside the range expected under lineage-label permutations. Under the primary model, the observed coefficient was −0.087 compared with a null mean of −0.001 (P = 0.001).

**S4.3 Alternative lineage graphs.** Because GMP has two parents in the lineage graph, we repeated the analysis after removing each incoming edge in turn. Estimates were similar to those obtained from the full graph (Table N4.2).

**S4.4 Close versus distant cell type pairs.** Cell type pairs separated by at most two lineage steps shared more variants than more distant pairs (15.7-fold versus

**Table N3.3** Each active category against the other two active categories pooled: odds ratio [95% CI]; BH-adjusted q, fewer than five variants in the category not tested.

| cCRE class | Specific | Multiple | Shared |
|---|---|---|---|
| PLS | 0.64 [0.55, 0.74]; q = $1.0\times10^{-8}$ | 1.40 [1.21, 1.63]; q = $1.6\times10^{-5}$ | 7.67 [4.68, 12.57]; q = $2.3\times10^{-11}$ |
| pELS | 0.75 [0.66, 0.84]; q = $3.7\times10^{-6}$ | 1.32 [1.17, 1.48]; q = $1.1\times10^{-5}$ | 1.72 [0.91, 3.27]; q = 0.12 |
| dELS | 0.69 [0.64, 0.74]; q = $1.3\times10^{-24}$ | 1.47 [1.37, 1.57]; q = $8.3\times10^{-26}$ | 0.69 [0.44, 1.08]; q = 0.12 |
| CA-H3K4me3 | 0.99 [0.75, 1.31]; q = 1.0 | 0.99 [0.75, 1.31]; q = 1.0 | – |
| CA-CTCF | 1.23 [0.98, 1.55]; q = 0.10 | 0.82 [0.65, 1.03]; q = 0.12 | – |
| CA-TF | 1.28 [0.90, 1.81]; q = 0.21 | 0.80 [0.56, 1.14]; q = 0.27 | – |
| CA | 1.60 [1.30, 1.98]; q = $1.4\times10^{-5}$ | 0.64 [0.52, 0.79]; q = $3.8\times10^{-5}$ | – |
| TF | 3.72 [1.81, 7.62]; q = $1.3\times10^{-4}$ | 0.28 [0.13, 0.56]; q = $2.1\times10^{-4}$ | – |
| No cCRE | 2.36 [2.14, 2.61]; q = $1.5\times10^{-67}$ | 0.43 [0.39, 0.48]; q = $2.4\times10^{-63}$ | – |

**Table N4.1** Distance coefficient under each expected count, with and without adjustment for background similarity.

| Expected count | Covariate | $\beta$ per step | 95% CI | Per-step ratio |
|---|---|---|---|---|
| Panel-wide rates | none | −0.182 | [−0.217, −0.147] | 0.833 |
| Within-pair (primary) | none | −0.087 | [−0.122, −0.052] | 0.917 |
| Panel-wide rates | background similarity | −0.053 | [−0.078, −0.027] | 0.949 |
| Within-pair | background similarity | +0.048 | [+0.027, +0.069] | 1.049 |

**Table N4.2** Distance coefficient after removing either edge into GMP.

| Graph | Within-pair $\beta$ per step [95% CI] | Per-step ratio | Panel-wide $\beta$ per step [95% CI] |
|---|---|---|---|
| Full | −0.087 [−0.122, −0.052] | 0.917 | −0.182 [−0.217, −0.147] |
| Without LMPP→GMP | −0.075 [−0.106, −0.044] | 0.927 | −0.152 [−0.185, −0.120] |
| Without CMP→GMP | −0.080 [−0.113, −0.047] | 0.923 | −0.174 [−0.206, −0.142] |

12.8-fold enrichment relative to expectation). Excess sharing declined monotonically across all six distance classes (16.0, 15.5, 13.8, 12.8, 11.0, and 10.4).

**S4.5 Direction of effect.** Among 11,416 instances where a variant was active in both members of a parent-child cell-type pair, only 48 (0.42%) showed opposite directions of effect. Sign discordance rates for individual edges are shown in Table N4.3.

**Table N4.3** Sign discordance on each parent–child edge.

| Edge | Active in both | Discordant | Rate [95% CI] |
|---|---|---|---|
| HSC→MPP | 1,435 | 0 | 0% [0, 0.17%] |
| MPP→LMPP | 1,097 | 2 | 0.18% [0.04, 0.58%] |
| MPP→CMP | 1,329 | 1 | 0.08% [0.01, 0.35%] |
| CMP→GMP | 1,230 | 1 | 0.08% [0.01, 0.38%] |
| CMP→MEP | 1,300 | 1 | 0.08% [0.01, 0.36%] |
| LMPP→CLP | 540 | 3 | 0.56% [0.16, 1.5%] |
| LMPP→GMP | 1,350 | 0 | 0% [0, 0.19%] |
| GMP→Mono | 661 | 1 | 0.15% [0.02, 0.70%] |
| MEP→Ery | 369 | 4 | 1.1% [0.37, 2.6%] |
| MEP→Mega | 609 | 8 | 1.3% [0.62, 2.5%] |
| CLP→B | 490 | 5 | 1.0% [0.39, 2.2%] |
| CLP→CD4 T | 348 | 6 | 1.7% [0.72, 3.5%] |
| CLP→CD8 T | 308 | 5 | 1.6% [0.62, 3.5%] |
| CLP→NK | 350 | 11 | 3.1% [1.7, 5.4%] |

### Supplementary Note S5: Agreement between ChromBPNet and fine-mapping

**S5.1 Enrichment of fine-mapped variants among ChromBPNet nomination.** The enrichment of fine-mapped variants among ChromBPNet-nominated variants was similar whether calculated across all accessible variants or restricted to variants carrying a posterior probability (Table N5.1). This indicates that the association between ChromBPNet nomination and fine-mapping support is not driven by the choice of analysis universe. The unadjusted odds ratio among the 36,200 accessible variants was 4.11 (95% CI 3.54–4.77), matching the value reported in the main text.

**Table N5.1** Enrichment of PIP $\geq$ 0.5 among active variants in two settings.

| Setting | Active, PIP $\geq$ 0.5 | Control, PIP $\geq$ 0.5 | Rate ratio | MH OR (95% CI) | Tarone P |
|---|---|---|---|---|---|
| All accessible (242,307) | 239 / 13,608 (1.756%) | 854 / 228,699 (0.373%) | 4.70 | 3.39 (2.92–3.94) | 0.66 |
| Carries a posterior (36,200) | 239 / 2,479 (9.641%) | 854 / 33,721 (2.533%) | 3.81 | 3.36 (2.87–3.92) | 0.73 |

**S5.2 Alternative nomination rules.** We compared the empirical-P activity definition with a fixed effect-size threshold ($|\text{logFC}| \geq 0.5$) and a magnitude-only rule selecting the top-ranked variants by $|\text{logFC}|$. All three rules produced similar enrichments for fine-mapped variants (Table N5.2).

When directly compared, the empirical-P and size-matched magnitude-only rules identified nearly identical numbers of fine-mapped variants among variants uniquely nominated by one rule or the other (Table N5.3).

**S5.3 Agreement across posterior-probability bins.** The fraction of variants nominated by ChromBPNet increased steadily with fine-mapping confidence

**Table N5.2** Matched-control enrichment for three calling rules.

| Rule (variants called) | Endpoint | Universe | Rate called / control | MH OR (95% CI) | Tarone P |
|---|---|---|---|---|---|
| Empirical $P$ (13,608) | PP $\geq$ 0.001 | All accessible | 18.2% / 14.7% | 1.10 (1.05–1.15) | 0.059 |
| Empirical $P$ (13,608) | PIP $\geq$ 0.5 | All accessible | 1.76% / 0.37% | 3.39 (2.92–3.94) | 0.66 |
| Empirical $P$ (2,479) | PIP $\geq$ 0.5 | Has posterior | 9.64% / 2.53% | 3.36 (2.87–3.92) | 0.73 |
| $\lvert$logFC$\rvert$ $\geq$ 0.5 (5,918) | PP $\geq$ 0.001 | All accessible | 19.4% / 14.8% | 1.21 (1.13–1.29) | 0.14 |
| $\lvert$logFC$\rvert$ $\geq$ 0.5 (5,918) | PIP $\geq$ 0.5 | All accessible | 2.42% / 0.40% | 4.41 (3.67–5.29) | 0.26 |
| $\lvert$logFC$\rvert$ $\geq$ 0.5 (1,149) | PIP $\geq$ 0.5 | Has posterior | 12.45% / 2.71% | 4.11 (3.39–4.99) | 0.37 |
| Size-matched (13,608) | PP $\geq$ 0.001 | All accessible | 18.2% / 14.7% | 1.11 (1.06–1.17) | 0.028 |
| Size-matched (13,608) | PIP $\geq$ 0.5 | All accessible | 1.75% / 0.37% | 3.43 (2.95–3.99) | 0.51 |
| Size-matched (2,480) | PIP $\geq$ 0.5 | Has posterior | 9.60% / 2.54% | 3.37 (2.88–3.94) | 0.72 |

**Table N5.3** Head-to-head comparison of the two size-matched rules.

| Endpoint | Empirical-$P$ only | Magnitude only | Exact binomial P |
|---|---|---|---|
| PP $\geq$ 0.001 | 288 (16.93%) | 289 (16.99%) | 1.0 |
| PIP $\geq$ 0.5 | 20 (1.18%) | 19 (1.12%) | 1.0 |

(Table N5.4). Among accessible variants, nomination rates increased from 5.6% in the lowest posterior-probability bin to 26.8% among variants with PIP $\geq$ 0.9.

**S5.4 All variants versus accessible variants.** The overlap between fine-mapping and ChromBPNet was stronger when analysis was restricted to accessible variants (Table N5.5). The difference reflects the large number of variants outside accessible chromatin that cannot be nominated by ChromBPNet.

The full list of 88 high-confidence variants is provided in Additional file 2; Table S1.

#### Supplementary Note S6: Molecular consequence, accessibility and cell type concordance of nominated variants

**S6.1 Consequence enrichment.** Selected consequence enrichments are shown in Table N6.1 (full results: Additional file 2: Table S6). Fine-mapping-only variants were enriched for coding and transcript-associated annotations, including missense, splice-region, and untranslated-region variants. In contrast, ChromBPNet-only variants were enriched for regulatory-region annotations and showed no enrichment for missense variation. Variants nominated by both methods showed the strongest enrichment for regulatory-region annotations.

**Table N5.4** Model-nominated fraction by posterior bin.

| Bin | All variants | Model nominated | % (95% CI) | Accessible variants | Model nominated | % (95% CI) |
|---|---|---|---|---|---|---|
| 0–0.01 | 105,595 | 1,016 | 0.96 (0.91–1.02) | 18,146 | 1,016 | 5.60 (5.27–5.94) |
| 0.01–0.1 | 73,237 | 943 | 1.29 (1.21–1.37) | 14,085 | 943 | 6.70 (6.29–7.12) |
| 0.1–0.5 | 11,980 | 281 | 2.35 (2.09–2.63) | 2,876 | 281 | 9.77 (8.74–10.91) |
| 0.5–0.9 | 1,610 | 78 | 4.84 (3.90–6.01) | 493 | 78 | 15.82 (12.87–19.31) |
| 0.9–1 | 1,474 | 161 | 10.92 (9.43–12.62) | 600 | 161 | 26.83 (23.44–30.52) |

**Table N5.5** Nomination classes among all and accessible variants.

| Class | All variants | Accessible variants |
|---|---|---|
| Both | 239 | 239 |
| Fine-mapping-only | 2,845 | 854 |
| Model-only | 2,240 | 2,240 |
| Neither | 188,572 | 32,867 |
| OR (95% CI) | 7.07 (6.16–8.12), Fisher $P = 3.6 \times 10^{-109}$ | 4.11 (3.54–4.77), $P = 4.3 \times 10^{-60}$ |

**S6.2 Accessible versus genome-wide backgrounds.** Because ChromBPNet can only nominate variants in accessible chromatin, enrichment analyses were repeated using both all variants and the subset accessible in at least one cell type as the comparison background. Restricting the background to accessible variants substantially reduced regulatory region enrichments (e.g. model-only variants: OR 11.05 to 2.40; doubly nominated variants: OR 14.08 to 3.06), indicating that part of the genome-wide enrichment reflects accessibility itself. However, regulatory region enrichment remained clearly positive after restricting the background, suggesting that the observed signal cannot be explained by accessibility alone.

**S6.3 Active-allele quantile.** ChromBPNet-only variants occupied more accessible chromatin than fine-mapping-only variants (Table N6.2). The median active-allele quantile was 0.692 for ChromBPNet-only variants compared with 0.607 for fine-mapping-only variants. All pairwise comparisons among nomination classes were significant (Table N6.3).

**S6.4 Cell-type concordance.** Lineage definitions used for concordance analyses are shown in Table N6.4.

Model-only variants showed significant lineage concordance, with 39.9% of variant-trait pairs active in the expected lineage compared with 31.0% expected under permutation (Table N6.5). Variants nominated by both methods showed stronger concordance (73.3% observed versus 45.9% expected). Similar results were obtained using expanded lineage definitions that included upstream progenitor populations.

#### Supplementary Note S7: Predicted accessibility gain and CRISPRi-validated target genes

**S7.1 Results across alternative analysis specifications.** We evaluated 2,304 alternative analysis specifications spanning variant sets, aggregation methods, filtering criteria, and TSS distance thresholds. Of these, 2,193 produced converged models and 2,141 yielded interpretable confidence intervals. For the strongest-variant aggregation method, 78.3% of analysis specifications produced odds ratios greater than one (median 1.50; IQR 1.09-2.17). For the mean aggregation method, 81.0% produced odds ratios greater than one (median 1.55). In contrast, only 27.1% of specifications using the fraction of variants predicted to increase accessibility produced odds ratios greater than one (median 0.84). The strongest-variant and mean-effect aggregation methods therefore showed a generally positive, although variable, relationship with CRISPRi-validated target genes. Of the 2,193 converged models, 111 produced nominal $P <$

**Table N6.1** Selected consequence enrichments against both backgrounds.

| Group | Class | n in group | Background | OR (95% CI), accessible | OR (95% CI), all |
|---|---|---|---|---|---|
| Fine-mapping only | Missense | 88 / 854; 341 / 2,845 | 403 / 32,867; 1,492 / 188,572 | 9.25 (7.27–11.78) | 17.08 (15.08–19.33) |
| Fine-mapping only | 5′ UTR | 64 / 854; 80 / 2,845 | 1,063 / 32,867; 1,600 / 188,572 | 2.42 (1.87–3.15) | 3.38 (2.69–4.25) |
| Fine-mapping only | Intron | 585 / 854; 1,995 / 2,845 | 25,790 / 32,867; 149,683 / 188,572 | 0.60 (0.52–0.69) | 0.61 (0.56–0.66) |
| Fine-mapping only | Stop-gained | 6 / 2,845 | 18 / 188,572 | below minimum count | 22.14 (8.78–55.81) |
| Fine-mapping only | Splice region | 8 / 854; 25 / 2,845 | 129 / 32,867; 467 / 188,572 | 2.40 (1.17–4.92) | 3.57 (2.38–5.35) |
| Fine-mapping only | 3′ UTR | 44 / 854; 207 / 2,845 | 1,043 / 32,867; 5,493 / 188,572 | 1.66 (1.22–2.26) | 2.62 (2.26–3.02) |
| Model only | Regulatory region | 1,111 / 2,240 | 9,553 / 32,867; 15,426 / 188,572 | 2.40 (2.20–2.62) | 11.05 (10.15–12.02) |
| Model only | Missense | 21 / 2,240 | 403 / 32,867; 1,492 / 188,572 | 0.76 (0.49–1.18) | 1.19 (0.77–1.83) |
| Model only | Synonymous | 12 / 2,240 | 443 / 32,867 | 0.39 (0.22–0.70) | 0.64 (0.36–1.13) |
| Both | Regulatory region | 133 / 239 | 9,553 / 32,867; 15,426 / 188,572 | 3.06 (2.37–3.96) | 14.08 (10.91–18.19) |

**Table N6.2** Active-allele quantile by nomination class.

| Group | n | Median | IQR | 10th–90th percentile |
|---|---|---|---|---|
| Both | 239 | 0.769 | 0.631–0.877 | 0.476–0.933 |
| Model only | 2,240 | 0.692 | 0.505–0.836 | 0.330–0.919 |
| Fine-mapping only | 854 | 0.607 | 0.376–0.814 | 0.197–0.914 |
| Neither | 32,867 | 0.451 | 0.222–0.699 | 0.092–0.863 |

**Table N6.3** Pairwise comparisons of the active-allele quantile.

| Comparison | Median difference | Rank-biserial | q |
|---|---|---|---|
| Both vs neither | 0.317 | 0.544 | $2.9 \times 10^{-47}$ |
| Model-only vs neither | 0.241 | 0.395 | $7.7 \times 10^{-215}$ |
| Both vs fine-mapping-only | 0.162 | 0.323 | $3.0 \times 10^{-14}$ |
| Fine-mapping-only vs neither | 0.155 | 0.247 | $8.2 \times 10^{-35}$ |
| Model-only vs fine-mapping-only | 0.085 | 0.147 | $3.3 \times 10^{-10}$ |
| Both vs model-only | 0.077 | 0.200 | $3.5 \times 10^{-7}$ |

**Table N6.4** Lineage definitions for the concordance analysis.

| Lineage | Traits (n) | Trait codes | Strict cell types | Loose adds |
|---|---|---|---|---|
| Erythroid | 14 | hct, hgb, hlr, hlr_p, irf, mch, mchc, mcv, mrv, mscv, rbc, rdw_cv, ret, ret_p | Ery, MEP | CMP |
| Megakaryocytic | 4 | mpv, pct, pdw, plt | Mega, MEP | CMP |
| Myeloid | 8 | baso, baso_p, eo, eo_p, mono, mono_p, neut, neut_p | GMP, Mono | CMP |
| Lymphoid | 2 | lymph, lymph_p | CLP, B cell, CD4 T, CD8 T, NK | LMPP |

**Table N6.5** Cell type concordance of variant–trait pairs.

| Cell type set | Group | Pairs | Observed | Expected (95% null interval) | Enrichment | P |
|---|---|---|---|---|---|---|
| Strict | Both | 861 | 73.3% | 45.9% (40.3–51.7%) | 1.60 | $< 10^{-4}$ |
| Strict | Model only | 4,667 | 39.9% | 31.0% (29.2–32.8%) | 1.29 | $< 10^{-4}$ |
| Loose | Both | 861 | 77.6% | 53.8% (48.4–59.4%) | 1.44 | $< 10^{-4}$ |
| Loose | Model only | 4,667 | 47.4% | 39.3% (37.4–41.1%) | 1.21 | $< 10^{-4}$ |

0.05, compared with 110 expected by chance. Significant specifications are shown in Fig. S6.

**S7.2 Effect of distance from transcription start sites.** The positive association between predicted accessibility gain and CRISPRi validation tended to strengthen when analyses were restricted to cCREs farther from the nearest protein-coding transcription start site. For the strongest-variant aggregation method, the primary-analysis odds ratio increased from 1.73 at >1 kb to 3.90 at >10 kb. Similar patterns were observed across alternative analysis specifications. These analyses are descriptive because sample sizes decrease substantially at larger distance thresholds.

**S7.3 Primary analysis specification.** Results from the primary analysis specification and related aggregation methods are shown in Table N7.1. The strongest-variant aggregation method produced the largest and most consistent positive associations. In contrast, the fraction of variants predicted to increase accessibility showed little evidence of association with validated target genes.

**Table N7.1** Primary specification across aggregation rules and variant sets.

| Specification | OR (95% CI) | P |
|---|---|---|
| Pooled, strongest variant (n = 1,467; 205 positive) | 3.90 (1.33–11.50) | 0.013 |
| Pooled, mean effect | 3.53 (0.99–12.55) | 0.052 |
| Pooled, fraction opening | 0.86 (0.61–1.20) | 0.37 |
| GWAS only, strongest variant | 3.35 (0.88–12.77) | 0.076 |
| eQTL only, strongest variant | 2.23 (0.52–9.57) | 0.282 |

**S7.4 Statistical power.** Under the primary analysis specification, the minimum detectable odds ratio at 80% power was approximately 4.7. Detecting an odds ratio of 2 would require approximately 7,300 distal elements, roughly five times the size of the current dataset.

#### Supplementary Note S8: Transcription-factor motifs and CRISPRi-validated target genes

**S8.1 Motif instances.** Fi-NeMo identified 82,394 motif instances across the 9,205 K562 cCREs. Of these, 53,900 confident motif instances located entirely within cCRE boundaries were used in the primary analyses. Most cCREs (8,702 of 9,205) contained at least one confident motif instance.

**S8.2 Motif associations with validated target genes.** Association results for all motif labels and scoring schemes are provided in Additional file 2: Table S7. Under the primary presence-based analysis, eight motif labels differed significantly between cCREs with and without validated target genes (Table N8.1). CTCF showed the strongest and most consistent negative association across all motif-scoring approaches. NFE2/NRF2 also showed a negative association in pooled analyses but was investigated further because of its similarity to several related bZIP-family motifs.

**S8.3 Multivariable analyses.** To determine whether motif associations could be explained by other genomic features, we fit multivariable models adjusting for ABC score, chromatin accessibility, GC content, distance to the nearest transcription start site, sequence constraint, element length, and predicted accessibility. CTCF remained negatively associated with target-gene status after adjustment (Table N8.2; full results in Additional file 2: Tables S8 and S9). NFE2/NRF2 also remained negative in these pooled analyses.

**Table N8.1** Motifs that differ between cCREs with and without a validated target gene.

| Motif | Positives with motif | Negatives with motif | OR | q |
|---|---|---|---|---|
| CTCF_2 | 161 / 1,259 | 1,545 / 7,946 | 0.61 | $1.6 \times 10^{-7}$ |
| NFE_0 | 437 / 1,259 | 3,332 / 7,946 | 0.74 | $1.9 \times 10^{-5}$ |
| HSF_2 | 5 / 1,259 | 127 / 7,946 | 0.25 | $1.7 \times 10^{-3}$ |
| ZNF_74 | 358 / 1,259 | 1,804 / 7,946 | 1.35 | $1.2 \times 10^{-4}$ |
| KLF-SP_0 | 805 / 1,259 | 4,585 / 7,946 | 1.30 | $2.1 \times 10^{-4}$ |
| ZNF_375 | 32 / 1,259 | 98 / 7,946 | 2.09 | $3.6 \times 10^{-3}$ |
| TAF_0 | 32 / 1,259 | 108 / 7,946 | 1.89 | $1.2 \times 10^{-2}$ |
| NRF_0 | 149 / 1,259 | 731 / 7,946 | 1.32 | $1.4 \times 10^{-2}$ |

**Table N8.2** Multivariable coefficients for CTCF and NFE2/NRF2.

| Motif | Scheme | Full-data | Bootstrap median (95% CI) | Non-zero / negative |
|---|---|---|---|---|
| CTCF | presence | −0.279 | −0.281 (−0.356, −0.209) | 500/500; 100% |
| NFE2/NRF2 | presence | −0.250 | −0.251 (−0.319, −0.189) | 500/500; 100% |
| CTCF | importance | −0.317 | −0.321 (−0.412, −0.230) | 500/500; 100% |
| NFE2/NRF2 | importance | −0.312 | −0.315 (−0.419, −0.235) | 500/500; 100% |

**S8.4 Sensitivity to study of origin.** To determine whether motif associations were driven by differences among CRISPRi studies, we repeated the analyses after excluding the Reilly dataset and within studies containing both positive and negative elements (Table N8.3; per-study counts in Additional file 2: Table S10). CTCF remained negatively associated with validated target genes across all analyses. In contrast, the NFE2/NRF2 association was substantially weakened after accounting for study of origin and was no longer supported in within-study comparisons.

**Table N8.3** CTCF and NFE2/NRF2 associations in source-sensitivity analyses.

| Analysis | Motif | OR (95% CI) | P |
|---|---|---|---|
| Excluding Reilly (1,183 positives) | CTCF | 0.62 (0.52–0.75) | $7.5 \times 10^{-8}$ |
| Excluding Reilly | NFE2/NRF2 | 0.75 (0.66–0.85) | $1.2 \times 10^{-5}$ |
| Crude, same 8,927 elements | CTCF | 0.57 | n/a |
| Mantel–Haenszel (four studies) | CTCF | 0.64 (0.53–0.79) | $1.7 \times 10^{-5}$ |
| Crude, same 8,927 elements | NFE2/NRF2 | 0.78 | n/a |
| Mantel–Haenszel (four studies) | NFE2/NRF2 | 0.95 (0.82–1.10) | 0.50 |

**S8.5 Motif identities.** The CTCF signal originated from a motif whose strongest TomTom matches were all CTCF-family motifs. In contrast, the NFE2/NRF2 signal arose from motifs with substantial similarity to several related bZIP transcription-factor families, including AP-1 and BATF. We therefore interpret the CTCF association as specific but treat the NFE2/NRF2 association as a broader bZIP-family signal.

#### Supplementary Note S9: Validation of the CTCF motif calls against K562 CTCF ChIP-seq

**S9.1 Validation of CTCF motif calls.** We used ENCODE K562 CTCF ChIP-seq data (ENCODE experiment ENCSR000DWE) to evaluate the CTCF motifs identified by Fi-NeMo. Of the 1,842 confident CTCF motif instances, 75.2% overlapped a K562 CTCF ChIP-seq peak. When the same motifs were randomly repositioned within their own cCREs, only 39.5% overlapped a peak. CTCF motif instances identified by Fi-NeMo were therefore enriched 1.9-fold relative to this within-element null ($P < 0.001$). The enrichment was specific to confident CTCF motifs: overlap rates were substantially lower for other motif classes (8.1%) and for low-confidence CTCF motif calls (4.2%). Most overlapping CTCF motifs lay near ChIP-seq summits, with a median distance of 8 bp. At the cCRE level, 77.6% of cCREs containing a CTCF motif overlapped a CTCF peak compared with 5.1% of cCREs lacking a CTCF motif (OR 64.6). Together, these results indicate that the Fi-NeMo CTCF calls correspond closely to experimentally observed CTCF binding in K562 cells.

**S9.2 CTCF motifs and validated target genes.** Consistent with the primary analyses, cCREs with a validated target gene were less likely to contain a CTCF motif than those without one (OR 0.61, 95% CI 0.51-0.72). This association remained after accounting for study of origin, distance to the nearest transcription start site, or both. The depletion was strongest among ChIP-supported CTCF motifs (study-adjusted OR 0.58, 95% CI 0.45-0.73), whereas motifs without ChIP support showed a weaker and non-significant association (study-adjusted OR 0.85, 95% CI 0.60-1.21). Similar estimates were observed among cCREs more than 10 kb from a transcription start site. As a comparison, the NFE2/NRF2 motif group showed a negative association in pooled analyses but not after adjustment for study of origin, consistent with the conclusion in Note S8 that this signal reflects differences among CRISPRi datasets rather than a robust motif-specific effect.

#### Supplementary Figures

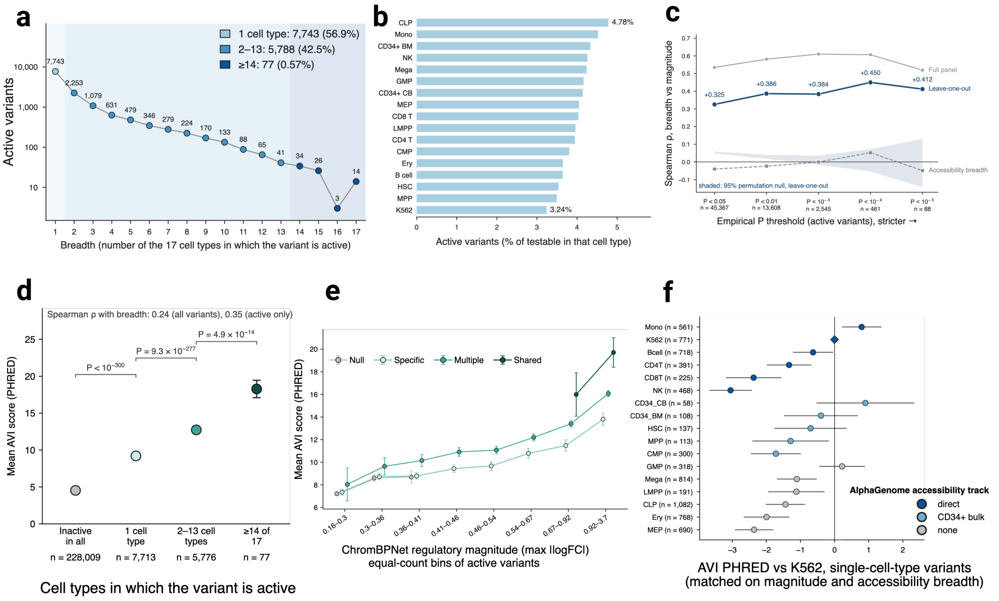

**Fig. S1 Breadth of predicted effects and comparison with AlphaGenome Variant Impact (AVI). a,** Distribution of regulatory breadth among active variants. Breadth is the number of the 17 cell types in which a variant is active (accessible with empirical $P < 0.01$). Shading indicates the three active breadth categories used throughout the manuscript: specific (1 cell type), multiple (2-13 cell types), and shared (14-17 cell types). **b,** Percentage of testable variants classified as active in each cell type. Cell types are ordered by activity rate. **c,** Correlation between breadth and predicted effect size across activity thresholds. Three estimators are shown: the full-panel estimator, the leave-one-out estimator used in the main text, and accessibility breadth. The shaded region shows the 95% permutation interval for the leave-one-out estimator. **d,** Mean AVI PHRED score by breadth category. Error bars show 95% confidence intervals. PHRED values of 10 and 20 correspond to the top 10% and top 1% of scored variants genome-wide. **e,** AVI PHRED score across bins of ChromBPNet regulatory magnitude. Variants were grouped by breadth category and binned by regulatory magnitude. Error bars show 95% confidence intervals. **f,** AVI differences are not explained by AlphaGenome accessibility-track coverage. Left, matched differences in AVI among single-cell-type variants grouped by active cell type. Marker fill indicates whether the corresponding cell type is directly represented by an AlphaGenome accessibility track, represented by bulk $CD34^+$ samples, or lacks a matching accessibility track. Right, matched AVI differences between multiple and specific variants, shown separately for variants active in at least one directly profiled cell type and for variants active in none. Error bars show 95% confidence intervals.

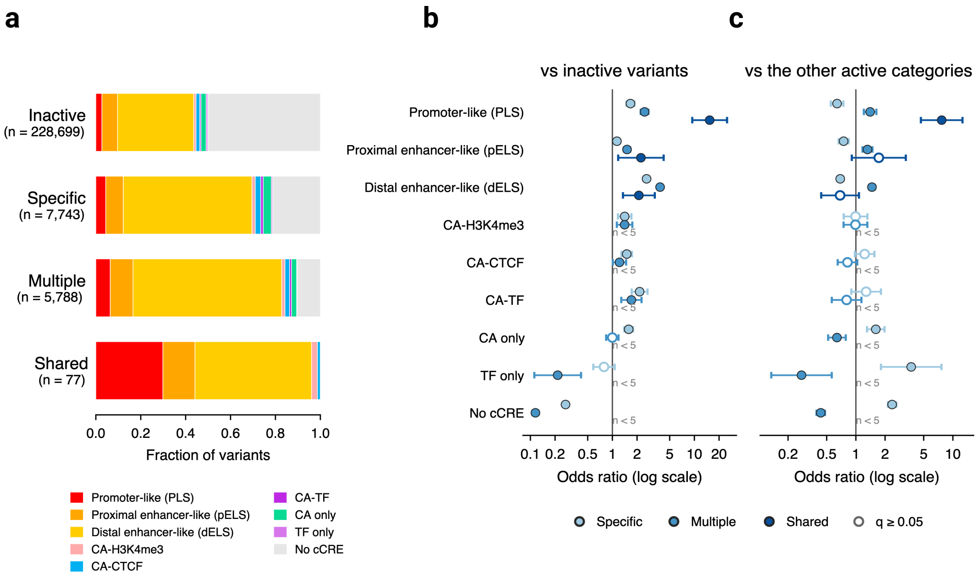

**Fig. S2 Regulatory element class and breadth of activity. a,** Fraction of variants in each ENCODE cCRE class (version 4, GRCh38) for the four breadth categories: inactive (0 cell types), specific (1), multiple (2–13), and shared (14–17), among the 242,307 variants accessible in at least one cell type. Colors follow the SCREEN cCRE classification scheme. **b,** Enrichment or depletion of each cCRE class in active variants relative to inactive variants, shown as odds ratios with 95% confidence intervals. Filled circles indicate $q < 0.05$ after Benjamini-Hochberg correction; open circles indicate $q \geq 0.05$. Classes with fewer than five variants were not tested. **c,** Enrichment or depletion of each active breadth category relative to the other two active categories combined. The composition of the active breadth categories differs significantly by cCRE class (permutation $P < 0.001$).

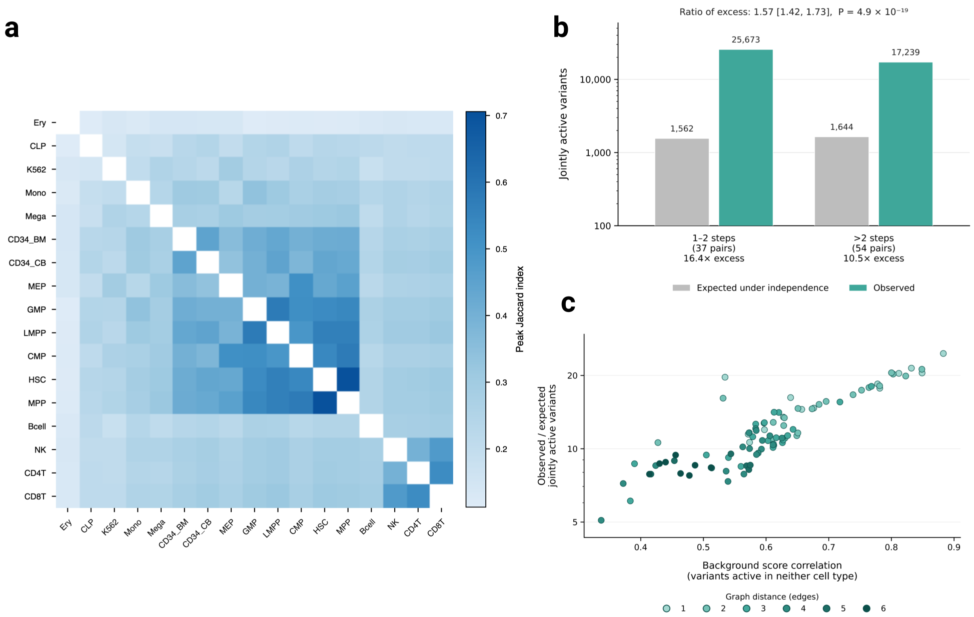

**Fig. S3 Accessibility similarity and co-activity across the hematopoietic lineage. a,** Pairwise similarity among all 17 cell types, measured as the Jaccard index of accessible peaks. **b,** Excess sharing of active variants among nearby (distance $\leq 2$) and more distant (distance $> 2$) cell-type pairs. Bars show observed counts of jointly active variants relative to expectations under independence using the panel-wide expectation. **c,** Excess sharing of active variants as a function of background similarity between cell types. Background similarity was calculated as the correlation of absolute ChromBPNet predictions among variants accessible in both cell types but active in neither. Points are colored by lineage distance. Cell type pairs with more similar background predictions also show greater excess sharing of active variants. **b,c** created in BioRender. Kazemi, M. (2026) https://BioRender.com/u2fjd32.

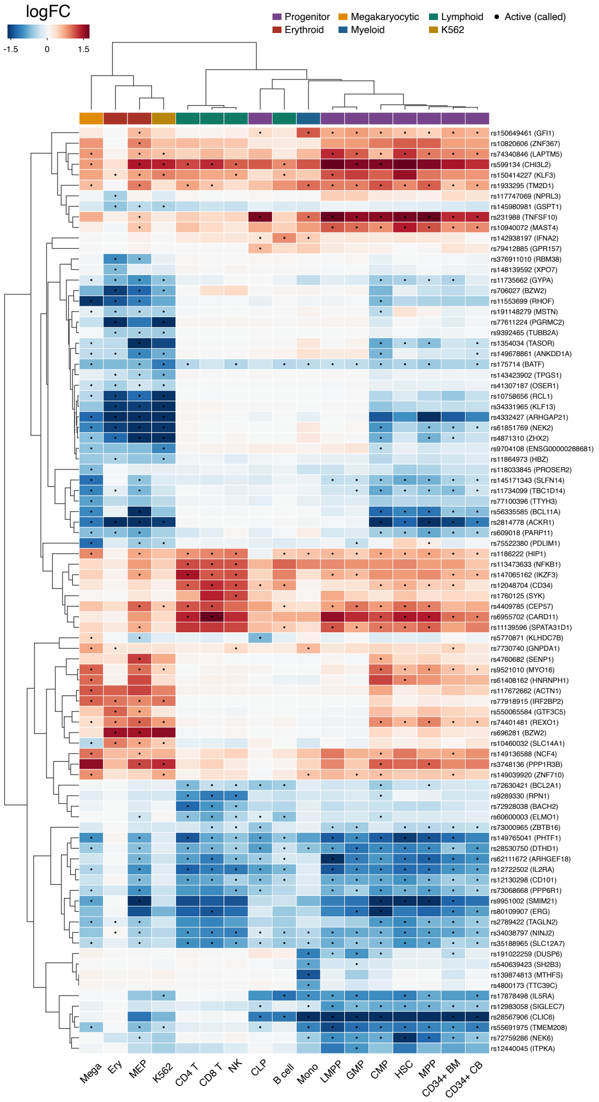

**Fig. S4 ChromBPNet predicted effects of the 88 high-confidence variants across 17 blood cell types.** High-confidence variants were defined as variants with PIP $\geq 0.5$ for at least one blood cell trait and empirical $P < 0.001$ in at least one accessible cell type. Rows represent variants (labeled by rsID and nearest protein-coding gene) and columns represent cell types. Colors show the predicted log fold change in accessibility (logFC), with positive effects in red and negative effects in blue. Rows and columns were clustered by correlation. The bar above the columns indicates lineage assignment. Dots mark variant-cell-type pairs classified as active (empirical $P < 0.01$). Nearest genes are reported for reference and are not target-gene assignments.

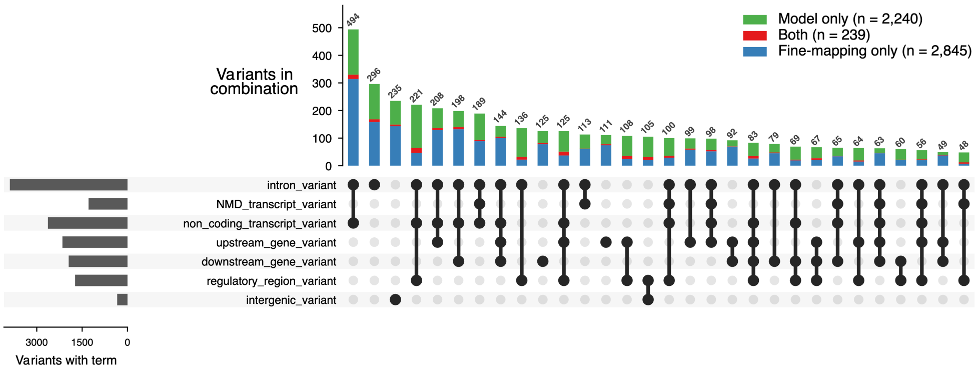

**Fig. S5 UpSet plot of VEP consequence combinations across variant nomination classes.** Variants were classified as model-only (active in at least one cell type and PIP $< 0.5$; n $= 2{,}240$), both (active and PIP $\geq 0.5$; n $= 239$), or fine-mapping-only (not active and PIP $\geq 0.5$; n $= 2{,}845$). Consequences correspond to Ensembl VEP annotations across all overlapping transcripts and regulatory features. Each variant was assigned to a single consequence combination representing its complete set of annotations. Horizontal bars show the number of variants carrying each consequence term. Vertical bars show the number of variants in each consequence combination, split by nomination class. The 30 most common combinations are shown, representing 3,900 of 5,324 variants (73.3%).

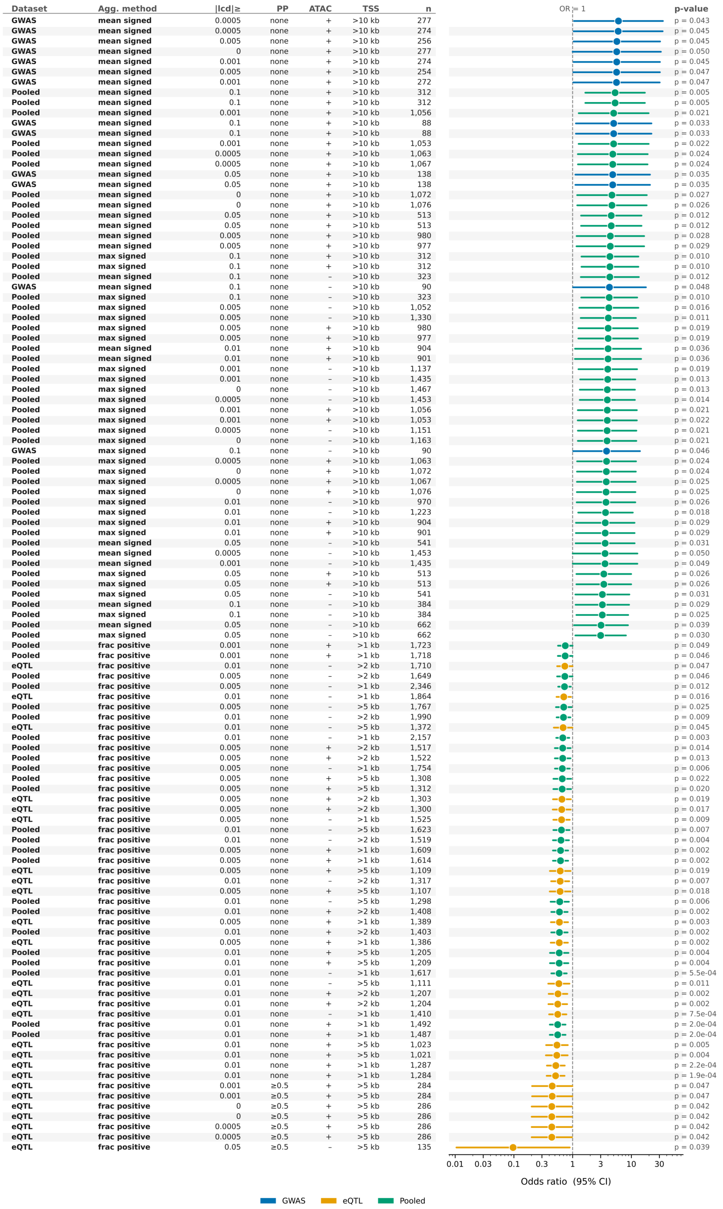

| Dataset | Agg. method | \|lcd\|≥ | PP | ATAC | TSS | n | p-value |
|---|---|---|---|---|---|---|---|
| **GWAS** | **mean signed** | 0.0005 | none | + | >10 kb | 277 | p = 0.043 |
| **GWAS** | **mean signed** | 0.0005 | none | + | >10 kb | 274 | p = 0.045 |
| **GWAS** | **mean signed** | 0.005 | none | + | >10 kb | 256 | p = 0.045 |
| **GWAS** | **mean signed** | 0 | none | + | >10 kb | 277 | p = 0.050 |
| **GWAS** | **mean signed** | 0.001 | none | + | >10 kb | 274 | p = 0.045 |
| **GWAS** | **mean signed** | 0.005 | none | + | >10 kb | 254 | p = 0.047 |
| **GWAS** | **mean signed** | 0.001 | none | + | >10 kb | 272 | p = 0.047 |
| **Pooled** | **mean signed** | 0.1 | none | + | >10 kb | 312 | p = 0.005 |
| **Pooled** | **mean signed** | 0.1 | none | + | >10 kb | 312 | p = 0.005 |
| **Pooled** | **mean signed** | 0.001 | none | + | >10 kb | 1,056 | p = 0.021 |
| **GWAS** | **mean signed** | 0.1 | none | + | >10 kb | 88 | p = 0.033 |
| **GWAS** | **mean signed** | 0.1 | none | + | >10 kb | 88 | p = 0.033 |
| **Pooled** | **mean signed** | 0.001 | none | + | >10 kb | 1,053 | p = 0.022 |
| **Pooled** | **mean signed** | 0.0005 | none | + | >10 kb | 1,063 | p = 0.024 |
| **Pooled** | **mean signed** | 0.0005 | none | + | >10 kb | 1,067 | p = 0.024 |
| **GWAS** | **mean signed** | 0.05 | none | + | >10 kb | 138 | p = 0.035 |
| **GWAS** | **mean signed** | 0.05 | none | + | >10 kb | 138 | p = 0.035 |
| **Pooled** | **mean signed** | 0 | none | + | >10 kb | 1,072 | p = 0.027 |
| **Pooled** | **mean signed** | 0 | none | + | >10 kb | 1,076 | p = 0.026 |
| **Pooled** | **mean signed** | 0.05 | none | + | >10 kb | 513 | p = 0.012 |
| **Pooled** | **mean signed** | 0.05 | none | + | >10 kb | 513 | p = 0.012 |
| **Pooled** | **mean signed** | 0.005 | none | + | >10 kb | 980 | p = 0.028 |
| **Pooled** | **mean signed** | 0.005 | none | + | >10 kb | 977 | p = 0.029 |
| **Pooled** | **max signed** | 0.1 | none | + | >10 kb | 312 | p = 0.010 |
| **Pooled** | **max signed** | 0.1 | none | + | >10 kb | 312 | p = 0.010 |
| **Pooled** | **mean signed** | 0.1 | none | - | >10 kb | 323 | p = 0.012 |
| **GWAS** | **mean signed** | 0.1 | none | - | >10 kb | 90 | p = 0.048 |
| **Pooled** | **max signed** | 0.1 | none | - | >10 kb | 323 | p = 0.010 |
| **Pooled** | **max signed** | 0.005 | none | - | >10 kb | 1,052 | p = 0.016 |
| **Pooled** | **max signed** | 0.005 | none | - | >10 kb | 1,330 | p = 0.011 |
| **Pooled** | **max signed** | 0.005 | none | + | >10 kb | 980 | p = 0.019 |
| **Pooled** | **max signed** | 0.005 | none | + | >10 kb | 977 | p = 0.019 |
| **Pooled** | **mean signed** | 0.01 | none | + | >10 kb | 904 | p = 0.036 |
| **Pooled** | **mean signed** | 0.01 | none | + | >10 kb | 901 | p = 0.036 |
| **Pooled** | **max signed** | 0.001 | none | - | >10 kb | 1,137 | p = 0.019 |
| **Pooled** | **max signed** | 0.001 | none | - | >10 kb | 1,435 | p = 0.013 |
| **Pooled** | **max signed** | 0 | none | - | >10 kb | 1,467 | p = 0.013 |
| **Pooled** | **max signed** | 0.0005 | none | - | >10 kb | 1,453 | p = 0.014 |
| **Pooled** | **max signed** | 0.001 | none | + | >10 kb | 1,056 | p = 0.021 |
| **Pooled** | **max signed** | 0.001 | none | + | >10 kb | 1,053 | p = 0.022 |
| **Pooled** | **max signed** | 0.0005 | none | - | >10 kb | 1,151 | p = 0.021 |
| **Pooled** | **max signed** | 0 | none | - | >10 kb | 1,163 | p = 0.021 |
| **GWAS** | **max signed** | 0.1 | none | - | >10 kb | 90 | p = 0.046 |
| **Pooled** | **max signed** | 0.0005 | none | + | >10 kb | 1,063 | p = 0.024 |
| **Pooled** | **max signed** | 0 | none | + | >10 kb | 1,072 | p = 0.024 |
| **Pooled** | **max signed** | 0.0005 | none | + | >10 kb | 1,067 | p = 0.025 |
| **Pooled** | **max signed** | 0 | none | + | >10 kb | 1,076 | p = 0.025 |
| **Pooled** | **max signed** | 0.01 | none | - | >10 kb | 970 | p = 0.026 |
| **Pooled** | **max signed** | 0.01 | none | - | >10 kb | 1,223 | p = 0.018 |
| **Pooled** | **max signed** | 0.01 | none | + | >10 kb | 904 | p = 0.029 |
| **Pooled** | **max signed** | 0.01 | none | + | >10 kb | 901 | p = 0.029 |
| **Pooled** | **mean signed** | 0.05 | none | - | >10 kb | 541 | p = 0.031 |
| **Pooled** | **mean signed** | 0.0005 | none | - | >10 kb | 1,453 | p = 0.050 |
| **Pooled** | **mean signed** | 0.001 | none | - | >10 kb | 1,435 | p = 0.049 |
| **Pooled** | **max signed** | 0.05 | none | + | >10 kb | 513 | p = 0.026 |
| **Pooled** | **max signed** | 0.05 | none | + | >10 kb | 513 | p = 0.026 |
| **Pooled** | **max signed** | 0.05 | none | - | >10 kb | 541 | p = 0.031 |
| **Pooled** | **mean signed** | 0.1 | none | - | >10 kb | 384 | p = 0.029 |
| **Pooled** | **max signed** | 0.1 | none | - | >10 kb | 384 | p = 0.025 |
| **Pooled** | **mean signed** | 0.05 | none | - | >10 kb | 662 | p = 0.039 |
| **Pooled** | **max signed** | 0.05 | none | - | >10 kb | 662 | p = 0.030 |
| **Pooled** | **frac positive** | 0.001 | none | + | >1 kb | 1,723 | p = 0.049 |
| **Pooled** | **frac positive** | 0.001 | none | + | >1 kb | 1,718 | p = 0.046 |
| **eQTL** | **frac positive** | 0.01 | none | - | >2 kb | 1,710 | p = 0.047 |
| **Pooled** | **frac positive** | 0.005 | none | - | >2 kb | 1,649 | p = 0.046 |
| **Pooled** | **frac positive** | 0.005 | none | - | >1 kb | 2,346 | p = 0.012 |
| **eQTL** | **frac positive** | 0.01 | none | - | >1 kb | 1,864 | p = 0.016 |
| **Pooled** | **frac positive** | 0.005 | none | - | >5 kb | 1,767 | p = 0.025 |
| **Pooled** | **frac positive** | 0.01 | none | - | >2 kb | 1,990 | p = 0.009 |
| **eQTL** | **frac positive** | 0.01 | none | - | >5 kb | 1,372 | p = 0.045 |
| **Pooled** | **frac positive** | 0.01 | none | - | >1 kb | 2,157 | p = 0.003 |
| **Pooled** | **frac positive** | 0.005 | none | + | >2 kb | 1,517 | p = 0.014 |
| **Pooled** | **frac positive** | 0.005 | none | + | >2 kb | 1,522 | p = 0.013 |
| **Pooled** | **frac positive** | 0.005 | none | - | >1 kb | 1,754 | p = 0.006 |
| **Pooled** | **frac positive** | 0.005 | none | + | >5 kb | 1,308 | p = 0.022 |
| **Pooled** | **frac positive** | 0.005 | none | + | >5 kb | 1,312 | p = 0.020 |
| **eQTL** | **frac positive** | 0.005 | none | + | >2 kb | 1,303 | p = 0.019 |
| **eQTL** | **frac positive** | 0.005 | none | + | >2 kb | 1,300 | p = 0.017 |
| **eQTL** | **frac positive** | 0.005 | none | - | >1 kb | 1,525 | p = 0.009 |
| **Pooled** | **frac positive** | 0.01 | none | - | >5 kb | 1,623 | p = 0.007 |
| **Pooled** | **frac positive** | 0.01 | none | - | >2 kb | 1,519 | p = 0.004 |
| **Pooled** | **frac positive** | 0.005 | none | + | >1 kb | 1,609 | p = 0.002 |
| **Pooled** | **frac positive** | 0.005 | none | + | >1 kb | 1,614 | p = 0.002 |
| **eQTL** | **frac positive** | 0.005 | none | + | >5 kb | 1,109 | p = 0.019 |
| **eQTL** | **frac positive** | 0.01 | none | - | >2 kb | 1,317 | p = 0.007 |
| **eQTL** | **frac positive** | 0.005 | none | + | >5 kb | 1,107 | p = 0.018 |
| **Pooled** | **frac positive** | 0.01 | none | - | >5 kb | 1,298 | p = 0.006 |
| **Pooled** | **frac positive** | 0.01 | none | + | >2 kb | 1,408 | p = 0.002 |
| **eQTL** | **frac positive** | 0.005 | none | + | >1 kb | 1,389 | p = 0.003 |
| **Pooled** | **frac positive** | 0.01 | none | + | >2 kb | 1,403 | p = 0.002 |
| **eQTL** | **frac positive** | 0.005 | none | + | >1 kb | 1,386 | p = 0.002 |
| **Pooled** | **frac positive** | 0.01 | none | + | >5 kb | 1,205 | p = 0.004 |
| **Pooled** | **frac positive** | 0.01 | none | + | >5 kb | 1,209 | p = 0.004 |
| **Pooled** | **frac positive** | 0.01 | none | - | >1 kb | 1,617 | p = 5.5e-04 |
| **eQTL** | **frac positive** | 0.01 | none | - | >5 kb | 1,111 | p = 0.011 |
| **eQTL** | **frac positive** | 0.01 | none | + | >2 kb | 1,207 | p = 0.002 |
| **eQTL** | **frac positive** | 0.01 | none | + | >2 kb | 1,204 | p = 0.002 |
| **eQTL** | **frac positive** | 0.01 | none | - | >1 kb | 1,410 | p = 7.5e-04 |
| **Pooled** | **frac positive** | 0.01 | none | + | >1 kb | 1,492 | p = 2.0e-04 |
| **Pooled** | **frac positive** | 0.01 | none | + | >1 kb | 1,487 | p = 2.0e-04 |
| **eQTL** | **frac positive** | 0.01 | none | + | >5 kb | 1,023 | p = 0.005 |
| **eQTL** | **frac positive** | 0.01 | none | + | >5 kb | 1,021 | p = 0.004 |
| **eQTL** | **frac positive** | 0.01 | none | + | >1 kb | 1,287 | p = 2.2e-04 |
| **eQTL** | **frac positive** | 0.01 | none | + | >1 kb | 1,284 | p = 1.9e-04 |
| **eQTL** | **frac positive** | 0.001 | ≥0.5 | + | >5 kb | 284 | p = 0.047 |
| **eQTL** | **frac positive** | 0.001 | ≥0.5 | + | >5 kb | 284 | p = 0.047 |
| **eQTL** | **frac positive** | 0 | ≥0.5 | + | >5 kb | 286 | p = 0.042 |
| **eQTL** | **frac positive** | 0 | ≥0.5 | + | >5 kb | 286 | p = 0.042 |
| **eQTL** | **frac positive** | 0.0005 | ≥0.5 | + | >5 kb | 286 | p = 0.042 |
| **eQTL** | **frac positive** | 0.0005 | ≥0.5 | + | >5 kb | 286 | p = 0.042 |
| **eQTL** | **frac positive** | 0.05 | ≥0.5 | - | >5 kb | 135 | p = 0.039 |

**Fig. S6 Significant associations across alternative analysis specifications.** Forest plot of all analysis specifications yielding nominal significance ($P < 0.05$) across the GWAS, eQTL, and pooled variant sets ($n = 111$). The left panel lists the analysis specification for each model, including data source, aggregation method, effect-size threshold, posterior-probability filter, accessibility filter, and transcription start site distance threshold. The right panel shows odds ratios with 95% confidence intervals. Colors indicate the variant set used (GWAS, blue; eQTL, orange; pooled, green). The dashed line marks no association ($OR = 1$).

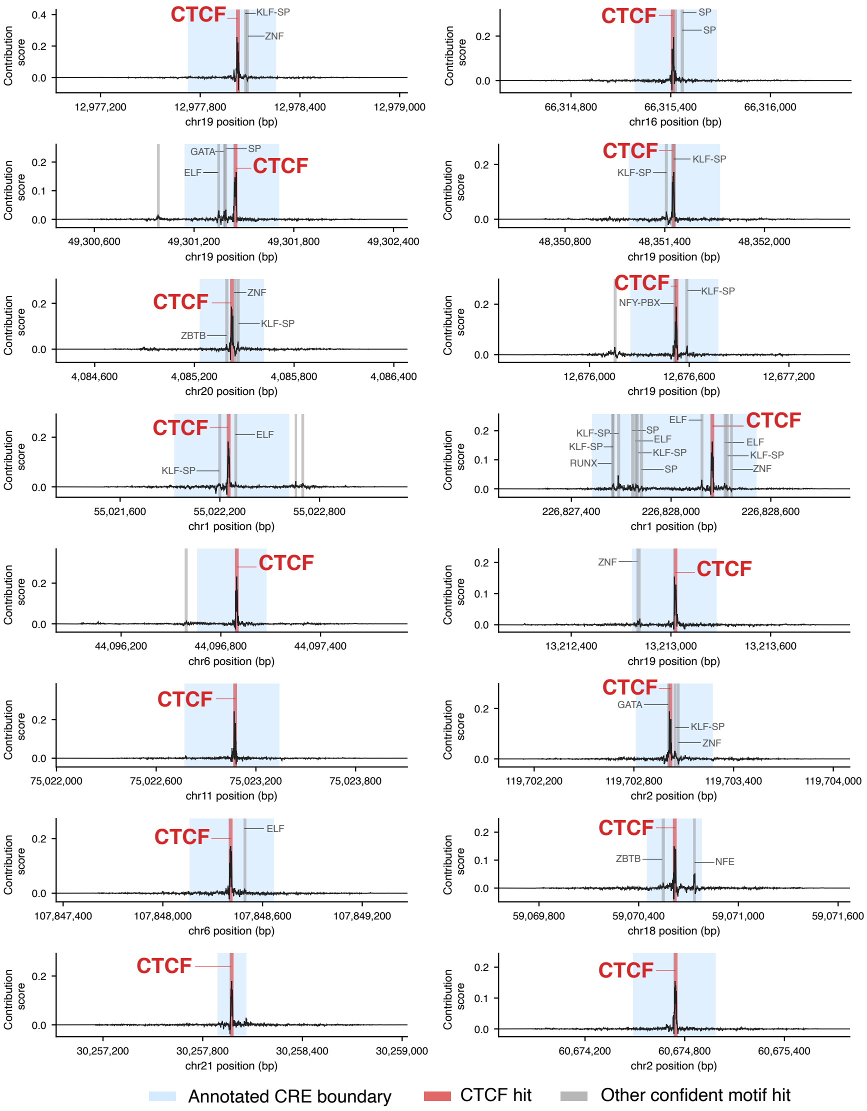

**Fig. S7 ChromBPNet contribution scores at CTCF motifs in K562 cCREs without validated target genes.** Sixteen K562 cCREs without a CRISPRi-validated target gene are shown, each containing a confident CTCF motif identified by Fi-NeMo. Panels are ordered by motif importance. Black lines show ChromBPNet contribution scores across the model input sequence. Light blue shading marks the annotated cCRE, the highlighted red region marks the CTCF motif instance, and gray regions mark other confident motif instances. Motif labels are shown above each panel, with CTCF highlighted in red. Confident motifs were defined as TOMTOM matches with $q < 0.05$. The y-axis is scaled independently for each panel.
